# An introgressed gene causes meiotic drive in *Neurospora sitophila*

**DOI:** 10.1101/2020.01.29.923946

**Authors:** Jesper Svedberg, Aaron A. Vogan, Nicholas A. Rhoades, Dilini Sarmarajeewa, David J. Jacobson, Martin Lascoux, Thomas M. Hammond, Hanna Johannesson

## Abstract

Meiotic drive elements cause their own preferential transmission following meiosis. In fungi this phenomenon takes the shape of spore killing, and in the filamentous ascomycete *Neurospora sitophila*, the *Sk-1* spore killer element is found in many natural populations. In this study, we identify the gene responsible for spore killing in *Sk-1* by generating both long and short-read genomic data and by using these data to perform a genome wide association test. Through molecular dissection, we show that a single 405 nucleotide long open reading frame generates a product that both acts as a poison capable of killing sibling spores and as an antidote that rescues spores that produce it. By phylogenetic analysis, we demonstrate that the gene is likely to have been introgressed from the closely related species *N. hispaniola*, and we identify three subclades of *N. sitophila*, one where *Sk-1* is fixed, another where *Sk-1* is absent, and a third where both killer and sensitive strain are found. Finally, we show that spore killing can be suppressed through an RNA interference based genome defense pathway known as meiotic silencing by unpaired DNA. *Spk-1* is not related to other known meiotic drive genes, and similar sequences are only found within *Neurospora*. These results shed new light on the diversity of genes capable of causing meiotic drive, their origin and evolution and their interaction with the host genome.

**Significance Statement:** In order to survive, most organisms have to deal with parasites. Such parasites can be other organisms, or sometimes, selfish genes found within the host genome itself. While much is known about parasitic organisms, the interaction with their hosts and their ability to spread within and between species, much less is known about selfish genes. We here identify a novel selfish “spore killer” gene in the fungus *Neurospora sitophila*. The gene appears to have evolved within the genus, but has entered the species through hybridization and introgression. We also show that the host can counteract the gene through RNA interference. These results shed new light on the diversity of selfish genes in terms of origin, evolution and host interactions.

## Introduction

Some genes can spread in a population even though they have negative effects on the organisms that carry them. These are often referred to as selfish genes and it is becoming increasingly clear that such genes can be important drivers of a large number of evolutionary patterns (Burt and Trivers, 2006; Rice, 2013; Werren, 2011). One class of selfish genetic elements are killer meiotic drivers (KMDs). When heterozygous at meiosis they can increase their own transmission by destroying or incapacitating meiotic products that do not carry them. KMDs have been found across eukaryotes, from plants to animals and fungi. In animals and plants they generally act in male meiosis by killing sperm. In fungi they kill sexual spores, and are therefore known as “spore killers” (Bravo Núñez et al., 2018; Lindholm et al., 2016).

A KMD must perform two tasks: kill non-carrier (sensitive) meiotic products, and ensure that it does not kill itself. They do this through either a poison-antidote mechanism, where they produce a poison that kills indiscriminately and an antidote that they keep to themselves, or through a killer-target mechanism, where the killer attacks a target element that can only be found in sensitive meiotic products. In order to successfully drive, the poison and antidote must always be inherited together and a target must never be inherited with a killer. For this reason many KMDs are found in regions of low recombination, such as inversions or on sex chromosomes (Larracuente and Presgraves, 2012; Lyon, 2003; Svedberg et al., 2018). Some KMDs only require a single gene to function, and so are not associated with regions of low recombination. This is often the case in fungi, where the majority of identified KMDs are single gene systems. In *Podospora*, a family of KMD genes named *Spok* are found at high frequencies (Grognet et al., 2014; Vogan et al., 2019). *Spok* genes produce a single protein that can act as both poison and antidote simultaneously, probably through the action of different functional domains (Vogan et al., 2019). *Podospora* also harbours *het-s*, a single-gene KMD which produces a prion that can cause drive by targeting the sensitive allele (Dalstra et al., 2003). In *Schizosaccharomyces pombe*, the large and highly diverse *wtf* gene family causes drive by producing poison and antidote products from overlapping transcripts generated from two different start codons (Hu et al., 2017; Nuckolls et al., 2017). Strains of *S. pombe* vary in *wtf* content and a single strain can carry up to 14 different driving *wtf* genes, generating extensive sexual incompatibilities between even closely related isolates (Bravo-Núñez et al., 2019; Eickbush et al., 2019).

As exemplified by the *wtf* genes, meiotic drive can often reduce the fertility of the organism. This puts the KMD in conflict with the rest of the genome, and suppressors are therefore expected to evolve. Suppression of drive has been observed in a number of species, but only in a few cases has the underlying mechanism of suppression been identified. For instance, in *D. simulans* RNA interference (RNAi) is necessary for suppression of a sex ratio KMD (Lin et al., 2018). The reduction of fertility is also hypothesized to drive reproductive isolation between populations, and some KMDs act as hybrid incompatibility loci (Frank, 1991; Hurst and Pomiankowski, 1991; Johnson, 2010; Phadnis and Orr, 2009). At the same time, if KMDs can cross reproductive barriers, they may be able to quickly establish themselves in sensitive populations (Meiklejohn et al., 2018; Sweigart et al., 2019).

In the ascomycete fungus *Neurospora*, three different spore killer KMDs have been identified (Turner and Perkins, 1979). *Sk-2* and *Sk-3* are found in *N. intermedia* and are both poison-antidote systems located in large, highly rearranged, non-recombining regions (Campbell and Turner, 1987; Svedberg et al., 2018). The *rsk* gene performs the antidote function for both *Sk-2* and *Sk-3*, but the gene performing the poison function in *Sk-2* (*rfk-1*) is not found in *Sk-3* (Hammond et al., 2012; Rhoades et al., 2019; Svedberg et al., 2018), meaning that an unknown gene must be responsible.

*Sk-1* is found in *N. sitophila* and was the first spore killer to be identified in *Neurospora*. In heterozygous *Sk-1* x sensitive crosses, nearly all spores carrying the sensitive genotype are killed, generating a distinct pattern of four black, viable spores, and four white, dead spores (Figure 1) (Turner and Perkins, 1979). Little is known about the genetics of *Sk-1* and its relationship to *Sk-2* and *Sk-3*, but an extensive sampling effort has identified and isolated over a hundred strains carrying *Sk-1*. While the *Sk-1* element is found globally and in approximately 15% of natural *N. sitophila* isolates, the variation in frequency among locations is large. The element appears to be absent in many populations, fixed in others, and in two well-sampled locations, Italy and Tahiti, both killers and sensitives are found in roughly equal frequencies (Jacobson et al., 2006; Turner, 2001).

**Figure 1:**
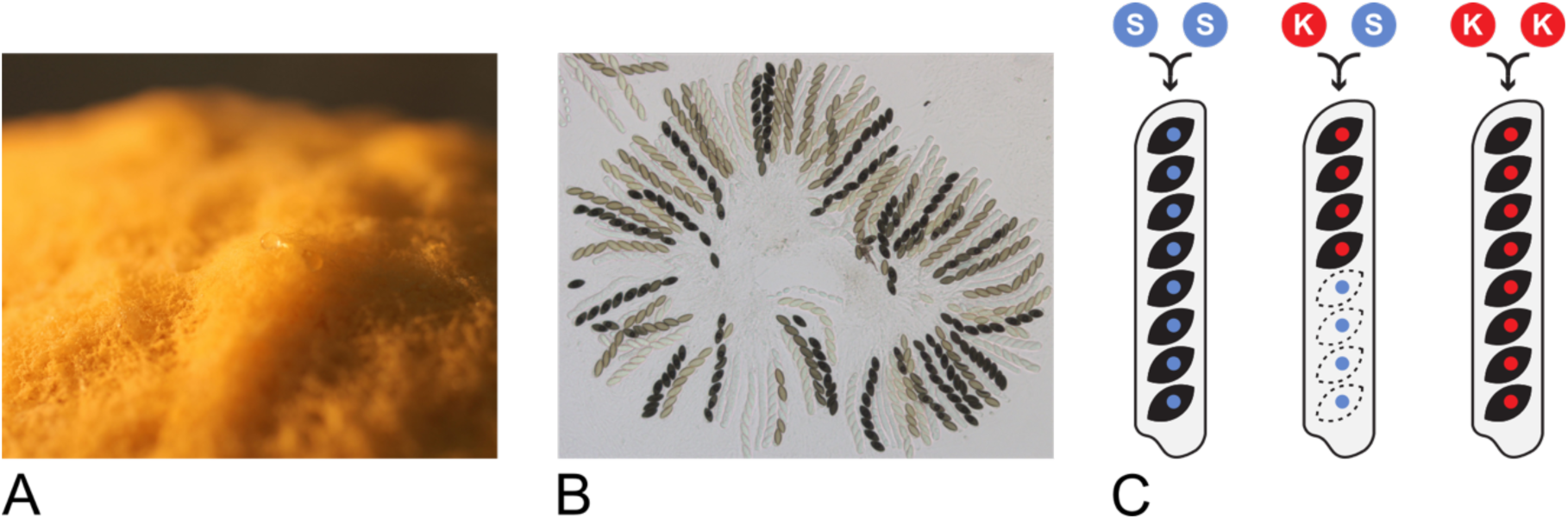
*Neurospora sitophila* and *Sk-1* spore killing. **(A)** *Neurospora sitophila* is closely related and morphologically similar to the model organism *N. crassa.* Both species produce large amounts of orange or yellow asexual spores (conidia). **(B)** Sexual ascospores in *N. sitophila*. *N. sitophila* is an obligate outcrosser and sexual reproduction takes place after a brief diploid stage when two haploid individuals encounter each other and cross. Each meiosis takes place in a cell known as an ascus and normally eight black, haploid ascospores are formed. **(C)** Schematic of produced ascospores in different crosses. In crosses which are homozygous for either the sensitive genotype (S-S) or the killer *Sk-1* genotype (K-K) eight black, viable ascospores are formed. In heterozygous crosses between sensitive and *Sk-1* killer strains (S-K), only four black, viable ascospores are formed. All surviving spores carry the *Sk-1* genotype and the sensitive spores degenerate during development and remain translucent. Spore killing occurs shortly after the cell walls surrounding the spores start to form. Homozygous *Sk-1* crosses produce eight spores as all spores carry a resistance factor which protects them from being killer.

In this study, we set out to identify the locus responsible for spore killing in *N. sitophila*. Using whole genome sequencing of 56 *N. sitophila* strains we show that *Sk-1* spore killing is caused by a single gene and demonstrate that it is responsible for both killing and resistance. Our data suggests that the *Sk-1* gene has been introgressed from a different *Neurospora* species, potentially *N. hispaniola*. Furthermore, we show that the population structure of *N. sitophila* is divided into three subclades, one of which is fixed for *Sk-1*, one where *Sk-1* is absent and a third where killers and sensitives intermix. Finally, we show that spore killing can be suppressed in certain crosses by an RNAi based genome defense mechanism known as meiotic silencing by unpaired DNA (MSUD).

## Results

### The locus responsible for spore killing is located on chromosome 6

We sequenced four *N. sitophila* strains (two *Sk-1* killers and two sensitives) using the PacBio RSII platform, generating high-quality genome assemblies, two of which had most chromosomes assembled from end to end (Table S1). Aligning these assemblies to the *N. crassa* OR74 genome assembly (Galagan et al., 2003) reveals that all four strains are largely collinear and that the two *Sk-1* strains do not carry any large rearrangements (Figure S1), unlike the *Sk-2* and *Sk-3* spore killer elements in *N. intermedia* (Svedberg et al., 2018).

We also generated short-read, paired-end sequencing data from 56 *N. sitophila* strains, using the Illumina HiSeq platform. Our sampling strategy aimed to capture the global diversity of *N. sitophila* and to include a balanced representation of both killers and sensitive strains (Figure S2, Table S2). Special attention was paid to two well-sampled populations in Tahiti (Turner, 2001) and Italy (Jacobson et al., 2006), which were both polymorphic for *Sk-1*.

In order to identify the locus responsible for spore killing, we conducted a genome wide association test. Illumina reads were mapped to the PacBio assembly of the killer strain W1434, SNPs were called and the association of each biallelic site to the killer phenotype was calculated using Fisher’s exact test (Figure 2A). Only 25 SNPs covering a 2 kbp region on Chromosome 6 showed a perfect association to the killer phenotype (Figure 2B, 2C). We call this region *sk1c1*, for *Sk-1 candidate 1*. A closer inspection revealed that in both killer and sensitive strains, the region contains a homolog to the *N. crassa* gene *NCU09865*. Sensitive strains carry the whole *NCU09865* sequence, but in the killer strains the allele is truncated and is missing the first half of the open reading frame; in its stead there is approximately 1 kbp of new sequence (Figure 2D). *NCU09865* is of unknown function in *N. crassa*, but it is predicted to contain a methyltransferase domain. This domain is present in the sensitive *N. sitophila* allele, but is partially missing in the *Sk-1* allele.

**Figure 2:**
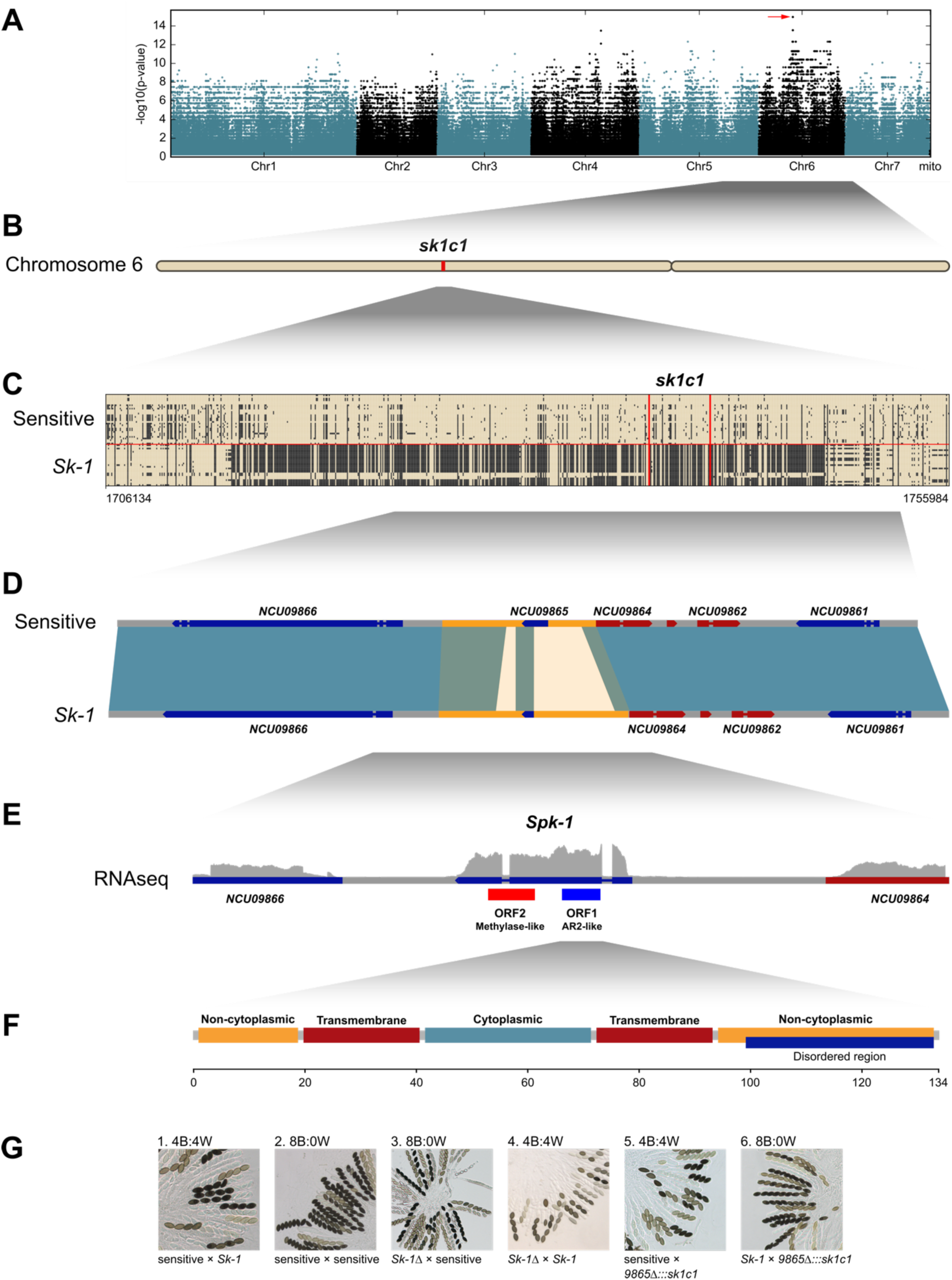
The locus responsible for spore killing. **(A)** Manhattan plot showing the association of 1,867,059 biallelic SNPs to the spore killer phenotype across the *N. sitophila* genome. A single locus (red arrow) on Chromosome 6 **(B)** shows complete association between genotype and phenotype. **(C)** The locus (*sk1c1*) contains 25 variable sites with complete association spanning 1933 bp and is contained within a highly diverged 23 kbp region which is found in 21 of 25 *Sk-1* strains. **(D)** In sensitive *N. sitophila* strains, *sk1c1* contains a homolog to the *N. crassa* gene *NCU09865*, but in *Sk-1* killer strains this gene is missing the first 375 bp, and it is instead replaced with 1.2 kbp of new sequence. **(E)** A transcriptomic analysis of a sensitive x *Sk-1* cross reveals a single transcript (*Spk-1*) in the *sk1c1^k^* region. The transcript contains two candidate open reading frames: ORF2 corresponding to the remaining part of *NCU09865* and ORF1 shows homology to *N. crassa* gene *NCU01957* (*AR2*). A molecular dissection of the transcript shows that only ORF1 is necessary for spore killing (Figure S5 and S6). **(F)** No functional protein domains could be identified in *Spk-1* ORF1, but an InterPro scan predict two transmembrane domains, a central cytoplasmic region and two flanking non-cytoplasmic region. **(G)** Images of asci from dissected perithecia. When crossing a sensitive strain to an *Sk-1* strain, 4 viable black and 4 dead white spores are produced, indicating spore killing (1). In a cross between two sensitive *N. sitophila* strains, 8 viable spores are formed in an ascus (2). When deleting the *sk1c1^k^* locus (*Sk-1*!) and crossing to a sensitive strain, killing is lost (3). When crossing *Sk-1*! to another *Sk-1* strain, spore killing is observed, indicating loss of resistance to spore killing (4). When inserting *sk1c1^k^* at the same locus in a sensitive strain (*9865Δ:::sk1c1*) and crossing it to a sensitive strain, killing is observed (5) and when crossed to *Sk-1*, killing is again observed (6). 5941 and W1432 were used as *Sk-1* tester strains, and 4739 and W1446 were used as sensitive tester strains.

### Confirmation of the killing and resistance activity of *sk1c1^k^*

We tested if the killer *sk1c1* allele (*sk1c1^k^*) is responsible for spore killing using three different strategies. First, we tested if it showed distorted segregation. A cross between a killer strain (W1446) and a sensitive strain (W1426) was performed and 46 sexual spores were picked and germinated separately. Segregation of the killer allele was assessed by PCR, with primers detecting a size polymorphism at *sk1c1*. All 46 spores carried the killer allele, showing complete segregation distortion.

We then generated a deletion mutant of the locus by deleting 2.8 kbp surrounding *sk1c1^k^* in strain W1434 (Figure S3A). When the deletion mutant was crossed to sensitive strains no killing was observed, and when it was crossed to killer strains killing was detected (Figure 2G). These results show that the *sk1c1^k^* is necessary both for killing sensitive strains and for resistance to killing from other *Sk-1* strains, and presumably itself.

Finally, we inserted *sk1c1^k^* at the same locus in a sensitive *N. sitophila* strain (W1426), by replacing *NCU09865*. When we crossed this knock-in strain to a sensitive strain (5941), spore killing was again detected (Figure 2G), and when we crossed it to an *Sk-1* strain (W1434), killing was not observed, indicating that *sk1c1^k^* is sufficient for both spore killing and for resistance. These results are consistent with a poison-antidote mechanism for *Sk-1*, where the presence of a resistance factor at *sk1c1^k^* can rescue any strain from killing, independent of genetic background.

### An ORF with homology to *NCU01957* is generating the causative agent for spore killing

We generated transcriptomic data from W1434 (*Sk-1*), 5940 (sensitive) and from a cross between the two strains. We identified a 1450 bp transcript consisting of three exons and two introns at *sk1c1^k^* in the data from W1434 and from the cross (Figure 2E). This result indicates expression of this transcript both during sexual development and vegetative growth. The two longest predicted ORFs are just over 400 bp, one of which corresponds to the *NCU09865* fragment (ORF2) and the other which shows homology to the *N. crassa* gene *NCU01957* (ORF1).

In order to determine which ORF causes spore killing or if both are necessary, we molecularly dissected the transcript. First, we performed RT-PCR using several primer pairs within the transcript, results of which support the existence of the 1450 bp locus spanning transcript with three exons and two introns (Figure S4). Second, we generated deletion mutants of each ORF (Figure S4, Figure S5). The deletion mutant of ORF1 showed both loss of killing when crossed to a sensitive strain and loss of resistance when crossed to a killer strain, matching the expected phenotype of a sensitive strain. The deletion mutant of ORF2 also showed loss of killing, but no loss of resistance. We then generated mutants where the start codon of each ORF was changed to a leucine (ATG to TTG) (Figure S6). Here we again observed both loss of killing and of resistance in the ORF1 mutant, but we observed loss of neither in the ORF2 mutant.

We interpret these results (summarized in Table S3) as showing that ORF1 is generating the causative agent behind spore killing in *Sk-1* strains, and that this single 405 bp ORF, translating into a predicted 134 aa protein, is both responsible for killing sensitive spores and for providing resistance against self-killing. We name this gene *Spk-1*.

### *Spk-1* has many different homologs in *Neurospora*

The SPK-1 protein contains no predicted functional domains, but an InterPro scan identifies two transmembrane domains, a central cytosolic region and two non-cytosolic flanks (Figure 2F). Searching the NCBI non-redundant protein database and FungiDB identifies no similar sequences outside of *Neurospora*, but several hits in *N. crassa* and *N. tetrasperma*. Two *N. crassa* genes show significant sequence identity (Figure S7). *NCU01957* (35 % amino acid identity) and *NCU16600* (32 % amino acid identity). *NCU01957* contains a Plasma Membrane ATPase domain not found in SPK-1, and a single amino acid mutation has been shown to cause sterility and produce empty asci (Randall and Metzenberg, 1998). No information on *NCU16600* function is available, but it is highly upregulated late in perithecial development, when ascospores start to be delimited (Wang et al., 2014). *Spk-1* also does not share any detectable homology with any of the other known spore killer genes in *Neurospora*, *Podospora* or *Schizosaccharomyces*.

We searched genome assemblies for 31 *Neurospora* strains from 7 different species (*N. crassa*, *N. discreta*, *N. hispaniola*, *N. intermedia*, *N. metzenbergii*, *N. sitophila* and *N. tetrasperma*, Table S4) for homologous sequences of *Spk-1* using tblastn. 115 hits over 100 aa were identified (including *NCU01957* and *NCU16600*) with amino acid sequence identities ranging from 24% to 98.5% (Figure S8). It is not clear to what extent these hits represent actual genes, but in most cases they correspond to potential ORFs of approximately equal or greater length than *Spk-1*.

### *sk1c1^k^* shows signals of introgression from *N. hispaniola*

The homologous sequence that shows the highest sequence identity (98.5% amino acid identity) to *Spk-1* is found in *N. hispaniola* strain 10403 (Villalta et al., 2009). This sequence is not only highly similar to *Spk-1*, but it also truncates *NCU09865* at the same site (Figure 3A, Figure S9). A phylogenetic analysis of the *NCU09865* fragment present in 10403 and the *Sk-1* killers, including alleles from several different species of *Neurospora* (Figure 3B), reveals that all *Sk-1* killers group with 10403, but are clearly divergent from all sensitive *N. sitophila* alleles.

**Figure 3:**
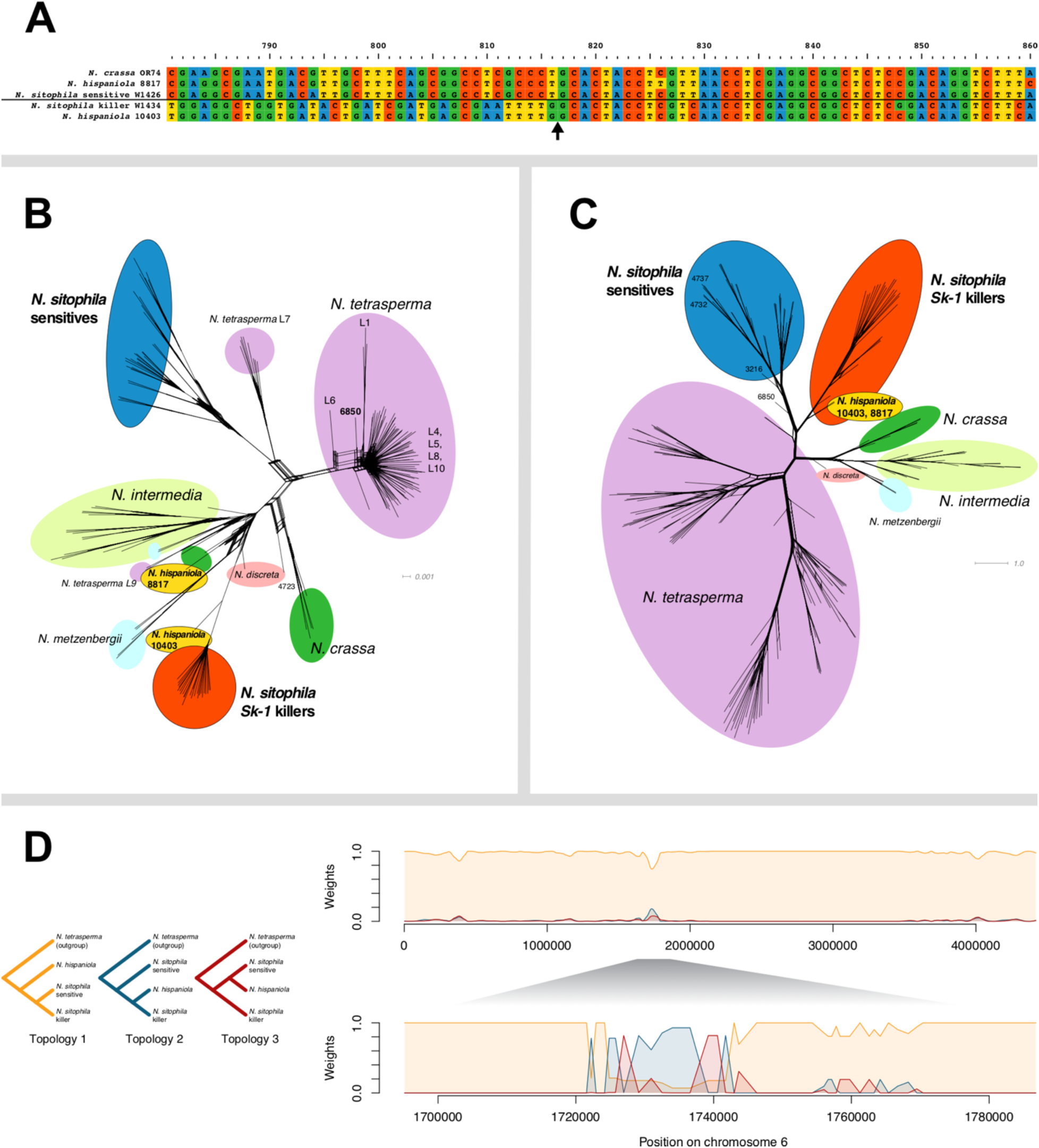
*Spk-1* is introgressed into *N. sitophila* from *N. hispaniola*. **(A)** Alignment of *NCU09865* segment. *N. crassa* OR74, *N. hispaniola* 8817 and sensitive *N. sitophila* W1426 all carry the complete gene, but *Sk-1 N. sitophila* W1434 and *N. hispaniola* 10403 are both missing the first 375 bp of the coding region. The black arrow marks the shared breakpoint of the deletion. **(B)** Phylogenetic network of the *NCU09865* fragment remaining in *Spk-1*. *N. sitophila Sk-1* and sensitives form two distinct clades. *N. hispaniola* 10403 groups with *Sk-1*, whereas N. hispaniola 8817 groups with neither. **(C)** Phylogenetic network of the concatenated sequences of the two genes flanking *NCU09865*: *NCU09864* and *NCU09866*. *Sk-1* and sensitive *N. sitophila* still form distinct clades, but here both *N. hispaniola* strains group with *Sk-1*. **(D)** Twisst plots showing signals of introgression from *N. hispaniola* into either *Sk-1* or sensitive strains of *N. sitophila.* The plot is generated by inferring phylogenetic trees in sliding windows containing 50 variable sites, then sampling subtrees with a strain from each clade from the tree and finally calculating the fraction of all subtrees that support each topology. The left panel shows the three possible phylogenetic topologies, the top panel shows distribution of topologies across chromosome 6 and the bottom panel the region surrounding *sk1c1*. In the *sk1c1* region, the dominant topology groups *Sk-1* strains with *N. hispaniola*, which is consistent with an introgressive scenario.

The two genes flanking *Spk-1* are located in the larger region that is fixed in 21 out of 25 killer strains and a phylogenetic analysis of these reveal that the *Sk-1* alleles are also most similar to alleles found in *N. hispaniola* (Figure 3C). These sequences cluster not only with sequences from strain 10403, but also with *N. hispaniola* strain 8817, which does not carry *Spk-1* or the truncated *NCU09865* sequence. The *N. sitophila* killers and the two *N. hispaniola* strains form a monophyletic group clearly separated from the sensitive *N. sitophila* strains. A phylogenetic analysis based on SNP data in sliding windows across the genome also identifies the *sk1c1^k^* allele as a candidate for introgression from *N. hispaniola* (Figure 3D), but on a genome wide level there is only evidence for very limited gene flow between the species (Figure S10 and Figure S11).

We interpret these results as suggesting that the killer locus has been introgressed into *N. sitophila* from *N. hispaniola*. However, we have a limited sample of *N. hispaniola* strains in our dataset and thus we cannot exclude the possibility that the source is another unknown species, or an unsampled and diverged population of *N. sitophila*.

### Three major clades are identified in *N. sitophila*

A phylogenetic analysis of whole-genome SNP data reveals clear population structure within *N. sitophila* and the investigated strains split into three major clades (Figure 4A), from now on referred to as Clade 1, 2 and 3. Clade 1 has a global distribution with strains from Asia, Oceania, Africa, the Caribbean, and Central and South America. Clade 2 contains strains from Asia, Africa and Europe and Clade 3 contains strains from Europe, Australia and continental United States. All strains from Tahiti, both *Sk-1* and sensitives, are placed in Clade 1, whereas all sensitive strains from Italy are placed in Clade 2 and all *Sk-1* strains from Italy are placed in Clade 3. In fact, Clade 2 is completely fixed for the sensitive genotype and Clade 3 for *Sk-1*. We only find evidence for limited gene flow between the three clades (Figure 4B), even though strains from Clade 2 and 3 were sampled at the same site and at the same time in Italy (Jacobson et al., 2006). These strains show no reduced fertility in laboratory crosses, but nevertheless, reproduction between strains of these clades appears to be rare in nature. Additionally, one strain of *N. sitophila* (6850) has been classified as resistant to *Sk-1* (Turner, 2001), but based on our analysis it shows comparatively strong differentiation from all other *N. sitophila* strains, and it may be classified as a different species.

**Figure 4:**
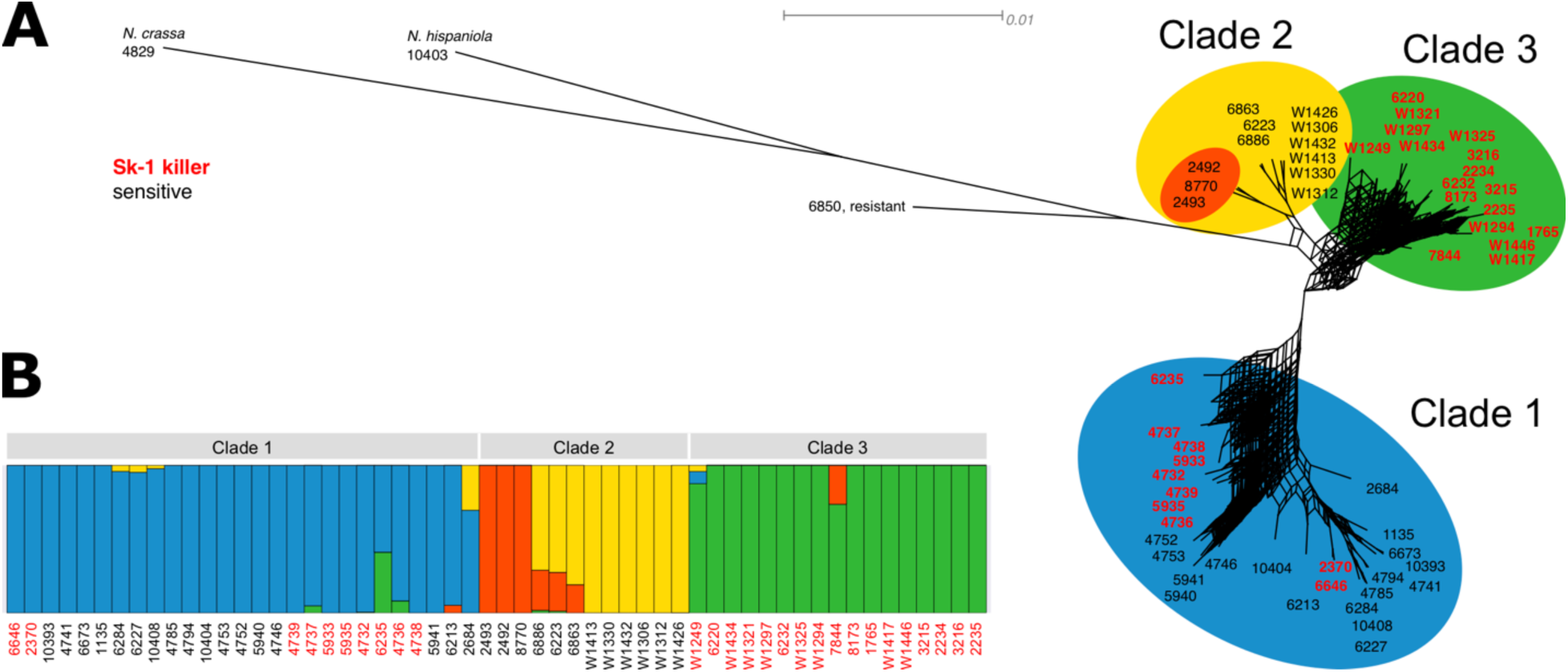
Population structure of *N. sitophila*. **(A)** A maximum likelihood phylogeny based on whole genome SNP data indicate that *N. sitophila* has clear population structure and can be split into three clades (Clade 1, 2 and 3). Both *Sk-1* and sensitive strains can be found in Clade 1, whereas Clade 2 is fixed for sensitives and Clade 3 is fixed for *Sk-1*, even though both Clade 2 and 3 can be found at the same sampling sites. **(B)** An admixture analysis of *N. sitophila* at K=4 further splits Clade 2 and shows limited signal for gene flow between the clades.

### Spore killing can be suppressed through RNAi

We wanted to investigate if the differences in frequency of *Sk-1* between the clades could be influenced by the genetic background. We therefore set out introgress *Sk-1* alleles from Italy and Tahiti into sensitive backgrounds from both regions and create isogenic lines by multiple rounds of backcrossing (Figure 5A). In the first round of crosses we observed normal levels of spore killing, but when backcrossing F1 individuals with a Tahiti sensitive parent to that sensitive parent, we saw a strong reduction in the number for four-spored asci (Figure 5B). Furthermore, after three generations of backcrossing, all lines generated from Tahitian sensitive parents had either completely lost spore killing or had a large reduction in spore killing (Figure S12). For the Italian sensitive lines, all lines displayed normal spore killing phenotypes. This pattern could be explained by *Spk-1* becoming unlinked from a modifying locus that improve killing efficiency, but when the Tahiti F1 individuals was outcrossed to another sensitive strain from either location, the number of four-spored asci was close to normal levels. This pattern suggests that the loss of spore killing is not caused by unlinking a modifier locus, but instead that it is an effect of increased relatedness between the killer and sensitive strains.

**Figure 5:**
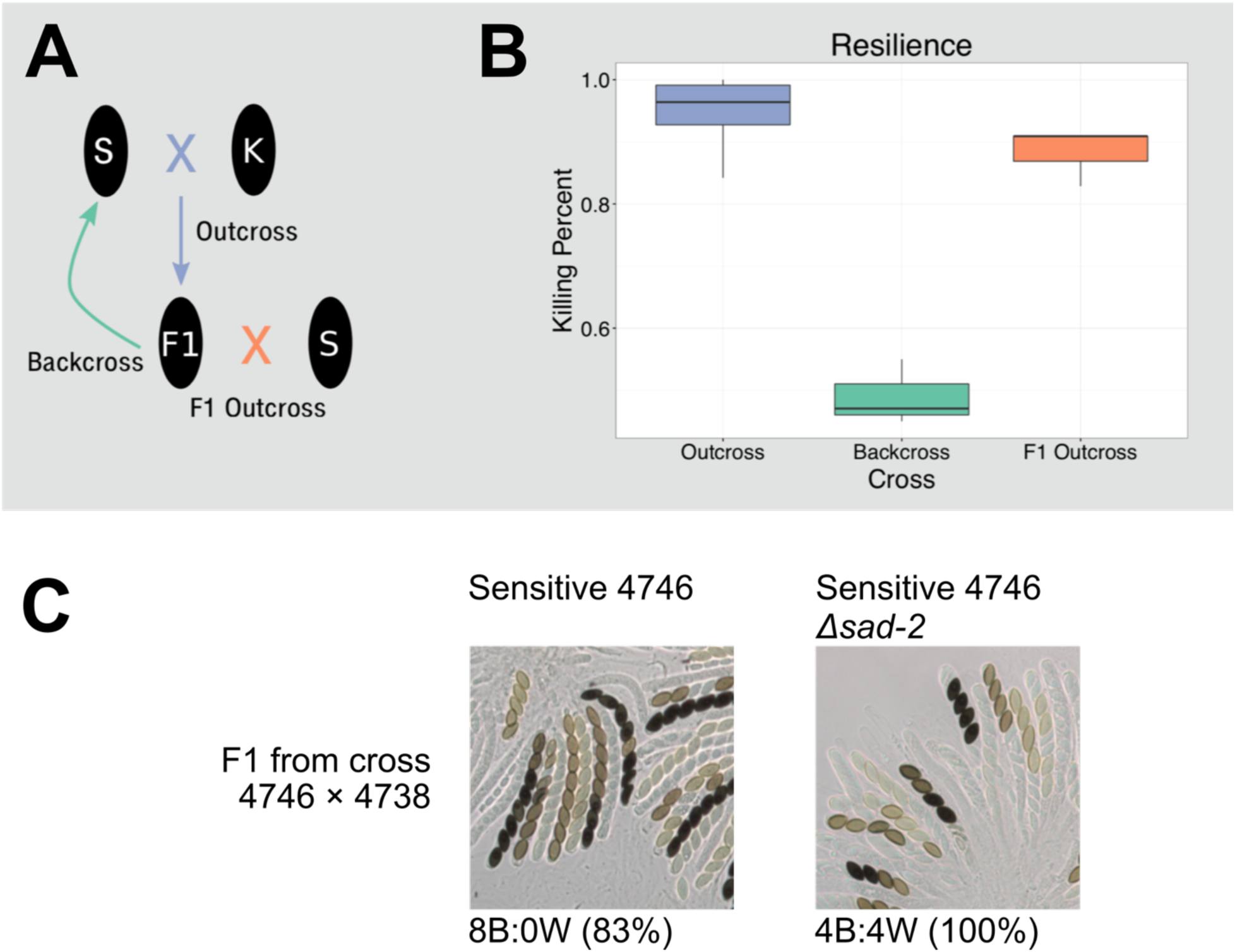
MSUD can suppress spore killing. **(A)** Crossing design to evaluate the strength of spore killing. S represents the sensitive parent 4746, K represents the *Sk-1* parent 4738. The F1 outcross was conducted to strain W1426. **(B)** Plot showing the fraction of asci exhibiting spore killing. **(C)** The effect of the *sad-2* deletion on the ability of backcrossed strains to kill. In a cross between a sensitive strain from Tahiti (4746) and a killer F1 from a cross between that strain and a killer strain (4738), killing is strongly reduced. Deleting the *sad-2* gene inactivates MSUD, and in a cross between 4746 *Δsad-2* and the killer F1, spore killing is restored to normal levels. This indicates that MSUD is responsible for suppressing spore killing in inbred crosses with a genetic background from Tahiti. This pattern is not seen if the sensitive strain comes from Italy.

MSUD is a genome defense mechanism identified in *Neurospora* that operates through the RNAi machinery to post-transcriptionally silence genes that are unpaired during meiosis (Shiu et al., 2001). Theoretically, MSUD should be capable of targeting and inactivating *Spk-1* as it is partially unpaired in a killer-sensitive cross, however MSUD activity has not been evaluated in *N. sitophila*. MSUD has been shown to have higher efficiency in inbred lines of *N. crassa* (Nagasowjanya et al., 2013), and we hypothesized that loss of killing could be driven by an increased MSUD activity caused by increased relatedness of the killer and sensitive strains. To test this hypothesis, we generated a strain that lacks functional MSUD by deleting *sad-2* (Shiu et al., 2006) in the sensitive Tahiti strain 4746. When *Sk-1* proficient offspring of 4746 were backcrossed to the *sad-2* deficient parent, close to 100% four spored asci were observed (Figure 5C, Figure S12, Figure S13), indicating that MSUD is indeed responsible for the loss of killing.

## Discussion

We have identified the gene responsible for the *Sk-1* spore killer phenotype in *N. sitophila*. It generates a single protein product of no more than 134 amino acids which is capable of both killing sensitive sibling spores and of protecting the spores producing the protein from self-killing. We have found no obvious mechanism behind either the killing or resistance phenotypes, but the gene carries predicted transmembrane domains, presenting the possibility that it could disrupt membrane integrity, as has been observed in some bacterial toxin-antitoxin systems (Lee and Lee, 2016; Unterholzner et al., 2013). Compared to the *wtf* genes in *Schizosaccharomyces* and the *Spok* genes in *Podospora*, *Spk-1* is significantly smaller (Grognet et al., 2014; Hu et al., 2017; Nuckolls et al., 2017; Vogan et al., 2019). The killing mechanism of the *wtfs* and *spoks* is still unknown, however the *wtf* genes also contain regions predicted to be transmembrane domain.

*Spk-1* shows no homology to any known spore killer gene, but appears to belong to a family of highly diverse *Neurospora* specific genomic sequences. It is still unclear to what extent these sequences are protein coding, but among them we found two annotated genes in the *N. crassa* reference genome. *NCU01957* is located very close to the mating type locus of *N. crassa*, but is not found at this location or elsewhere in most other investigated *Neurospora* genomes. Exceptions are *N. hispaniola*, where it is found at the same locus, and *N. sitophila*, where a highly similar gene is found at approximately 20 kbp distance. *NCU01957* is of unknown function, but a strain carrying a single amino acid mutation displayed both stunted vegetative growth and total sterility (Randall and Metzenberg, 1998). *NCU16600* is also of unknown function, but it is upregulated during ascosporogenesis (Wang et al., 2014). The highly diverse set of homologous sequences is reminiscent of the *wtf* genes in *Schizosaccharomyces*, with complex phylogenetic patterns and extensive presence-absence polymorphisms (Bravo-Núñez et al., 2019). Further work is necessary to determine if these are genes with a regular function for normal sexual development, or if they could be selfish genetic elements that have either gone to fixation or are controlled by some suppressing mechanism.

The *sk1c1^k^* allele found in *N. sitophila Sk-1* strains is highly diverged from sensitive *N. sitophila* strains and together with the flanking genes show greater similarity to the closely related species *N. hispaniola*. Strain 10403 carries a copy of *Spk-1* that shows both high sequence and structural similarity to the one found in *Sk-1* strains, suggesting that it may also be able to cause spore killing. Only four strains of *N. hispaniola* have ever been collected (Villalta et al., 2009), and we have not been able to observe spore killing when crossing these to each other or in hybrid crosses to *N. tetrasperma*. This could mean that *Spk-1* in 10403 is no longer functional, but it can also indicate that the strains available carry an unknown resistance factor and further sampling would be necessary to assess whether *Spk-1* is actively driving within the species.

While *sk1c1^k^* appears to be introgressed from *N. hispaniola* there is little sign of introgression elsewhere in the genome. *N. sitophila* and *N. hispaniola* show high levels of reproductive isolation with each other and we have so far not observed viable offspring in any crosses between the species. While selfish genetic elements such as meiotic drivers have often been argued to contribute to restricting gene flow between species (Frank, 1991; Hurst and Pomiankowski, 1991), here we see the opposite result, where meiotic drive instead may have facilitated a limited exchange of genetic material. Similar results have been observed in *D. simulans* and *D. mauritiania* where the Winters sex ratio element has crossed between species despite strong reproductive barriers (Meiklejohn et al., 2018). At this time we cannot exclude that *Sk-1* entered *N. sitophila* through a yet undiscovered species or population of *Neurospora* that shows greater sexual compatibility with both *N. sitophila* and *N. hispaniola*. We also do not know if *Spk-1* evolved in *N. hispaniola* or if it originated in another species and has spread within *Neurospora* through multiple introgressive steps.

Using whole genome SNP data we can show that *N. sitophila* is split into three clades which show remarkably different patterns of spore killing. While 15% of all *N. sitophila* strains carry the *Sk-1* element, these are not evenly distributed within the clades. Clade 3 is fixed for *Sk-1*, Clade 2 is fixed for sensitives, and Clade 1 contains both genotypes. This population structure is especially puzzling as strains from Clade 2 and 3 were sampled at the same location at the same time and show no signs of reduced fertility when crossed under laboratory conditions, but still show only very limited signatures of gene flow. Some mechanism must prevent individuals from these two clades to cross in nature, but unfortunately we currently know too little about the ecology and life cycle of *N. sitophila* to have a good understanding of what this might be.

In Clade 1, we find both *Sk-1* and sensitive strains. It is unclear if *Sk-1* has recently been introduced into the clade and is on its way to fixation or it if it is maintained as a stable polymorphism due to unknown fitness costs of carrying the spore killer element. One tantalizing suggestion is that the *Sk-1* polymorphism can be explained by the observed suppression through MSUD. While we only observe strong suppression in highly isogenic crosses, we do not know how common such crosses are in nature. If inbreeding is a common pattern in *N. sitophila*, and MSUD is more efficient between inbred strains, MSUD may be sufficient enough to keep *Sk-1* at a low frequency. It is also possible that MSUD is more efficient under natural growing conditions where environmental factors may promote a higher efficiency.

RNAi has been shown to be essential for suppressing sex linked meiotic drive in *D. simulans* and is suspected to be involved in other *Drosophila* species as well (Aravin et al., 2004; Ellison and Bachtrog, 2019; Lin et al., 2018). In *D. simulans* the sex ratio driver *Dox* is suppressed by two genes named *Nmy* and *Tmy* (Lin et al., 2018; Tao et al., 2007a, 2007b). Both genes are partial duplicates of *Dox* and generate small RNAs which silence *Dox* expression and prevents inactivation of Y bearing sperm. The production of small RNAs have also been observed from duplicated genes that are suspected to cause sex ratio distortion in *D. miranda* (Ellison and Bachtrog, 2019).

Here we show that the RNAi based MSUD system can act as a suppressor of spore killing in highly isogenic crosses. Transcriptional silencing through MSUD is not dependent on gene duplications, but can act directly on genes that are unpaired at meiosis. Silencing through gene duplication may actually not be possible in *Neurospora*, as duplicated sequences are targeted and mutated by the RIP (Repeat Induced Point-mutations) system, which will introduces high levels of C-to-T mutations in larger sequences found at least twice in a genome (Galagan and Selker, 2004). Both MSUD and RIP are thought to defend against selfish genetic elements such as transposable elements, and MSUD should at least in theory be capable of defending against invading meiotic drivers that are sufficiently divergent to not be able to pair at meiosis (Hammond, 2017). It is interesting to note that both *Spk-1* in *N. sitophila* and *rfk-1*, which is the causative factor behind the *Sk-2* spore killer element in *N. intermedia* (Rhoades et al., 2019), manage to evade MSUD in most crosses. In the case of *rfk-1*, the small size together with proximity to pairing DNA appears to protect it from being silenced (Rhoades et al., 2019) and in the case of *Spk-1*, a potentially higher MSUD efficiency in isogenic crosses is enough to tip the balance against the spore killer gene. We can hypothesize that MSUD is a sufficient defence against most KMDs which try to invade a population of *Neurospora*, and the only ones that are able to persist or go to fixation are the rare cases that have some feature that allows them to evade genome defense mechanisms such as MSUD.

Most of the killer meiotic drive elements that were initially discovered and which have been studied in greatest detail are large multi-gene systems found in regions of suppressed recombination (Larracuente and Presgraves, 2012; Lindholm et al., 2016; Lyon, 2003; Svedberg et al., 2018). It has sometimes even been stated that meiotic drive can only be caused by the interaction of multiple loci and that the association with regions of low recombination is a defining characteristic of the phenomenon. *Spk-1* joins a growing number of recently discovered single gene KMDs which contradict this view. In fungi, four out of five known meiotic drive gene families are caused by single genes which can cause drive on their own (Dalstra et al., 2003; Grognet et al., 2014; Nuckolls et al., 2017; Rhoades et al., 2019). All of these gene families appear to have evolved independently, suggesting that there is ample potential for *de novo* evolution of single gene KMDs, but if this is the case, why have we not found more of these systems? Single genes are less likely to be tightly linked to deleterious mutations, which could make them more likely to go to fixation and become invisible in crosses. They are also less likely to be tightly linked to visible phenotypes that would allow easy detection. Using genomic techniques to detect segregation distortion in controlled crosses, together with sampling more non-model systems has the potential to unmask a vast diversity of cryptic meiotic drive.

## Methods

### Strains

All natural isolates used in this study are listed in Table S2. Strains were ordered from the Fungal Genetics Stock Center (http://www.fgsc.net, (McCluskey et al., 2010)), except those whose names start with W (e.g., W1434), which were provided by D. Jacobson (Jacobson et al., 2006). Most strains were annotated as either *Sk-1* spore killers or sensitives, and we verified these phenotypes and tested the strains without annotation by crossing them to both *Sk-1* killer and sensitive tester strains carrying the *fluffy* mutation, and patterns of spore killing were then observed using a microscope. The tester strains are listed in Table S5. Strains used for molecular characterization are listed in Table S6.

Furthermore, we used the *N. crassa* OR74 genome (Galagan et al., 2003), corrected for assembly errors discovered by (Galazka et al., 2016), as well as a dataset of 92 *N. tetrasperma*, one *N. sitophila* and one *N. hispaniola* strain (Corcoran et al., 2016).

### Culture conditions

For maintenance, strains were grown in ambient conditions on Vogel’s medium (Vogel, 1956). Crosses were conducted in liquid synthetic crossing media (SC) with filter paper and no added sucrose. Two different protocols were used to isolated ascospores. 1. Single ascospores were collected from the sides of culture tubes and plated onto 2% water agar or Vogel’s medium. These plates were heat treated at 60°C for 60 minutes to induce germination and kill any conidia or mycelia that may have been transferred along with the ascospores. Germinating ascospores were excised from the water agar and transferred to Vogel’s media for growth and storage. Spore killing was evaluated at the point when ascospores could first be observed on the walls of the culture tubes. Mature perithecia were then removed and dissected. Rosettes of ascospores were imaged at 200x magnification on an Olympus BX50 Fluorescence microscope. 2. Ascospores were placed in 500 µL water, heat chocked for 30 minutes and then plated on Vogel’s medium. Spore killing was evaluated after 15 days. Rosettes were inaged on a a Leica DMBRE compound microscope.

### Genome sequencing

Whole genome short-read data was collected from 56 *N. sitophila* strains (Table S2). Conidia were inoculated into 50 ml plastic culture tubes containing 10 ml of 3 % liquid malt extract medium. The tubes were incubated at 30 °C on a rotary shaker for 2–3 days and were then harvested by removing the mycelium from the culture tubes which was squeezed between filter paper to remove excess liquid. It was then cut into pieces and approximately 100 µg was allotted into 1.5 ml Eppendorf tubes, which were stored at −20 °C until extraction. Genomic DNA was extracted using the Fungal/Bacterial Microprep Kit (Zymo, www.zymo.com) and sent to the SNP&SEQ Technology Unit (Science for Life Laboratory, Uppsala, Sweden), where libraries were prepared and sequenced on an Illumina HiSeq 2500 system.

Four *N. sitophila* strains (Table S1) were sequenced using the PacBio RSII platform (Pacific Biosciences). The strains were cultured by inoculating 200 ml of 3% liquid malt extract medium in 500 ml Erlenmeyer flasks with conidia. The flasks were placed on a rotary shaker at 30 °C for 3–4 days. The cultures were harvested by removing the mycelium from the liquid, placing it on a filter paper, which was folded between several layers of tissue paper and pressed to remove excess liquid. The mycelium was cut into smaller pieces and approximately 1 g was allotted into 2 ml tubes with screw-on caps. These tubes were then stored at −20 °C. To extract genomic DNA, two tubes of each strain were freeze-dried overnight and macerated using a TissueLyzer II bead-beater (Qiagen). Two 2 mm metal beads were placed in each tube, which were then shaken for 20 s at 25 Hz. If a tube contained large pieces of freeze-dried mycelium, it was shaken for another 10 s. This procedure was repeated until no large pieces remained. No tube was shaken for more than 40 s in total. DNA was finally extracted using Genomic Tip G-500 columns (Qiagen) and cleaned using the PowerClean DNA Clean-Up kit (MoBio Labs). Library preparation and sequencing was performed at the Uppsala Genome Center (Science for Life Laboratory, Uppsala, Sweden) using the C4 chemistry, P6 polymerase and four SMRT cells per strain (Pacific Biosciences).

### Genome assembly

Raw PacBio sequence data was filtered using the SMRT Analysis package and assembled *de novo* using the HGAP 3.0 assembler (Chin et al., 2013) (Pacific Biosciences, https://github.com/PacificBiosciences/). Sequencing information and assembly statistics is shown in Table S1. The W1426 and W1434 assemblies were also annotated using MAKER (Cantarel et al., 2008), using protein and transcript sequences from *N. crassa* OR74 (Galagan et al., 2003).

Raw Illumina HiSeq reads were assembled *de novo* using ABySS (Simpson et al., 2009), with the following parameters: Kmer size = 64, bubble size = 3, seed length=200. Sequencing information and assembly statistics is shown in Table S2.

### Whole genome alignment

Synteny and collinearity of the PacBio assemblies was investigated through whole genome alignment to the high-quality *N. crassa* OR74 reference genome (Galagan et al., 2003) and to each other. The PacBio assemblies were aligned using MUMmer (Kurtz et al., 2004) with the parameters “nucmer –c 200 –b 2000”. The alignments were visualised using MUMmer’s plotting function mummerplot. Chromosome numbers were also assigned based on the corresponding *N. crassa* chromosome.

### SNP calling

Raw Illumina HiSeq reads were cleaned from adapter contamination using CutAdapt (Martin, 2011) and trimmed using Trimmomatic (Bolger et al., 2014). The reads were then mapped to the W1434 PacBio assembly with BWA (Li and Durbin, 2009). The resulting BAM file was deduplicated with picard (2019), and complex regions were realigned with GATK IndelRealigner (McKenna et al., 2010). SNPs were then called with GATK (first using -T HaplotypeCaller -bamWriterType CALLED_HAPLOTYPES -stand_emit_conf 10.0 -stand_call_conf 20.0 -gt_mode DISCOVERY -emitRefConfidence BP_RESOLUTION, then all VCF files were merged using -T GenotypeGVCF -sample_ploidy 1 -includeNonVariantSites). Sites with missing data and variants from regions called as repetitive with RepeatMasker were removed using VCFtools (Danecek et al., 2011).

### Genome wide association of variation to *Sk-1*

In order to identify the locus or loci responsible for spore killing in *Sk-1* strains, the association of each SNP to the killing phenotype was calculated by a custom Python script which performed Fisher’s exact test for each variable site. The output was visualised with the python script manhattan-plot.py, which was downloaded from https://github.com/brentp/bio-playground/tree/master/plots/.

### Characterization of the killer locus

The gene located at *sk1c1* was identified as a homolog to *NCU09865* by the MAKER genome annotation package (Cantarel et al., 2008), when annotating the sensitive strain W1426. The similarity of the truncated fragment in the killer strain W1434 to *NCU09865* was confirmed using BLAST (Altschul et al., 1990). Homologous sequences to *NCU09865* found in other closely related species had in many cases been annotated as having a methylase or methyltransferase domain. We confirmed this in *NCU09865* by using EBI’s HMMer tool (Finn et al., 2015), which reports the presence of a Methyltransf_11 domain (PF08241.11).

### Generating and analyzing transcriptomic data

Transcriptomic data was generated from vegetative tissue of *Sk-1* strain W1434, sensitive strain 5940, and from a sexual cross between both strains. Each strain was grown separately on solid Vogel’s medium on petri dishes covered in cellophane for 2 days at 25 °C, under 12h:12h light–dark conditions. Hyphal tissue was then harvested from the surface of the cellophane with a sterile scalpel, and stored at −80°C. The sexual cross was performed on synthetic crossing medium (pH 6.5, 1.5% sucrose; (Westergaard and Mitchell, 1947)) with Wattman filter paper embedded into the medium, and covered with cellophane in 100 mm diameter petri dishes.

The harvested hyphal tissue were immediately frozen in liquid nitrogen and stored at 80°C until RNA extraction. Next, 150 mg of the frozen tissue was ground together with liquid nitrogen and total RNA was extracted using the RNeasy Plant Mini Kit (Qiagen, Hilden, Germany). RNA quality was checked on the Agilent 2100 Bioanalyzer (Agilent Technologies, USA). All RNA samples were treated with DNaseI (Thermo Scientific). And sequencing libraries were prepared using the NEBNext Ultra Directional RNA Library Prep Kit for Illumina (New England Biolabs). mRNA was selected by purifying polyA+ transcripts (NEBNext Poly(A) mRNA Magnetic Isolation Module, New England Biolabs). Finally, the three paired-end libraries were sequenced with Illumina HiSeq 2500 at the SNP and SEQ Technology platform, generating 125 bp paired-end reads. The raw reads were trimmed using CutAdapt and Trimmomatic. They were then mapped to PacBio assemblies of the parental strains using STAR (Dobin et al., 2013). In the case of the cross, the *sk1c1^k^* region from W1434 was added as a separate contig to the 5941 genome assembly. Transcripts were finally called using cufflinks.

### Phylogenomic analysis of *N. sitophila*

A whole genome phylogenetic analysis was performed with RAxML (Stamatakis, 2014) using the SNPs called against the W1434 PacBio assembly, which had been converted into a fasta file containing all sites with GATK VariantsToTable and a custom Python script. RAxML was run with the following parameters: raxmlHPC-HYBRID-AVX -m GTRCAT -# 100 -f a. Phylogenies of each chromosome were generated the same way and phylogenetic discordance was visualized by merging the trees into a Consensus Network with Splitstree (Huson and Bryant, 2006).

### Identification of SPK-1 homologs

We searched the NCBI non-redundant protein database and FungiDB’s (Stajich et al., 2012) protein database using blastp and default parameters. We also used tblastn (Gertz et al., 2006) to locate homologous sequences to the SPK-1 protein in 31 Neurospora genome assemblies, most of which were high-quality assemblies based on PacBio sequencing data (Table S4). Hits longer than 100 aa were extracted, flanking sequence corresponding to 60 aa were added. Candidate start and stop codons then were manually identified and the sequences were trimmed. We aligned these sequences using MAFFT 7.407 (Katoh and Standley, 2013) with default parameters and inferred a maximum likelihood phylogeny using IQ-TREE (Nguyen et al., 2015) with “parameters -st AA -m TEST -bb 1000 -wbt -alrt 1000”.

### Phylogenetic analysis of the killer locus

Phylogenies of the *NCU09865* fragment located at *sk1c1^k^* and two neighbouring genes were also inferred. Homologous sequences to the *NCU09865* fragment were extracted from *de novo* assemblies of all *N. sitophila* strains analyzed in this project, together with 29 further *Neurospora* assemblies from (Svedberg et al., 2018) and 92 *N. tetrasperma*, *1 N. sitophila* and *1 N. hispaniola* assembly from (Corcoran et al., 2016), by using genBlastG (She et al., 2011) to extract the top hit to the *N. crassa NCU09865* protein sequence from each genome assembly. The sequences were aligned with MAFFT (Katoh and Standley, 2013) and the alignment was manually trimmed using AliView (Larsson, 2014). Finally, phylogenetic relationships were inferred using RAxML, with the same parameters as above. The two neighbouring genes were extracted in the same way, by extracting the top hits to *NCU09864* and *NCU098766*. These two alignments were then concatenated and analyzed with RAxML as above.

### Detecting introgression from *N. hispaniola* across the genome

We used the program Twisst (Martin and Belleghem, 2017) to study how the phylogenetic signal varies across the genome, by following the pipeline described at https://github.com/simonhmartin/twisst. VCF files with SNPs called against W1434 were converted to the “.geno” file format used by Twisst, and local phylogenies were inferred in non-overlapping windows containing 50 SNPs using the Neighbour-joining algorithm of PhyML (Guindon et al., 2010), with the following parameters: “phyml_sliding_windows.py -w 50 --windType sites --model GTR --genoFormat haplo”. Twisst was then used to cluster local tree topologies with the parameters “run_twisst_parallel.py -T 4 --method complete”. Finally the output was plotted using the R script provided by the Twisst package.

### Admixture analysis

Admixture between the different *N. sitophila* populations and from *N. hispaniola* was assessed using the software ADMIXTURE (Alexander et al., 2009) using the W1434 SNP dataset and running the analysis from 2 to 8 ancestral populations.

### PCR verification of segregation of *sk1c1^k^*

Strains W1426 and W1446 were mated in liquid synthetic crossing media (SC) with filter paper and no added sucrose. Single ascospores were collected from the sides culture tubes and plated onto 2% water agar. These plates were heat treated at 60°C for 1 hour to induce germination and kill any conidia or mycelia that may have been transferred along with the ascospores. Germinating ascospores were excised from water agar plates and transferred to Vogel’s media from growth and storage. 46 progeny were obtained.

DNA was extracted using the Chelex 100 protocol (Bio-Rad). Primers were designed around a length polymorphism in the *sk1c1* region. The primers SK1C1F - ACCTCATCGTTCTGCAGCCCTCAT and SK1C1R - CTCCGAGCGAGGCTTGTGTGC produce a PCR product of ∼900bp in the parental killer strain W1446 and a product of ∼600bp in the parental sensitive strain W1426. The PCR protocol was as follows: initial denaturing at 98°C for 30s, 30 cycles of 98°C for 15s, 60°C for 30s, 72°C for 30s, and a final elongation step of 72°C for 10m. Products were visualized on a 1% agarose gel.

### Genetic transformation and strain construction

*N. sitophila* conidia were transformed by electroporation as described by Margolin et al (Margolin et al., 1997). Homokaryotic transformants were isolated with the method of Ebbole and Sachs (1990). Gene-deletion and transgene-insertion vectors were constructed by double joint PCR (DJ-PCR; Yu et al., 2004) using primers listed in Table S7. Left (L) and right (R) flanks were amplified from genomic DNA of the transformation host. The *hph* center fragment (C) of vectors v134, v178, v235, and v236 was amplified from pTH1256.1 (GenBank MH550659.1). The *Sk-1-hph* center fragment for v205a was amplified from pAY3.3. The Sk-1ORF1[ATG>TTG]-hph center fragment for v205b was amplified from pNR139.5. The Sk-1ORF2[ATG>TTG]-hph center fragment for v205c was amplified from pNR141.1.

Plasmids pAY3.3, pNR139.5, and pNR141.1 were constructed by amplifying the *Spk-1* gene from strain W1434 with primers 1507 (5′ TTTTGCGGCCGCAGCATTTTCACCTTGGCCGTGAG 3′) and 1508 (5′ TTTTGAATTCGATATGGGGAACGGGATTGTGGA 3′) and cloning the gene between the *Not*I and *Eco*RI sites of plasmid pTH1256.1. Site-directed mutagenesis of the *Spk-1* gene in pNR139.5 and pNR141.1 was performed essentially as described by the QuikChange II Site-Directed Mutagenesis Kit (Agilent Technologies). For pNR139.5, the predicted *Spk-1* ORF1 start codon was mutated to TTG using primers 1747 (5′ ATGTCGCAGATTGAACAACAC 3′) and 1748 (5′ TTACTATCATGTTAGAAAGGAG 3′). For pNR139.5, the predicted *Spk-1* ORF2 start codon was mutated to TTG using primers 1751 (5′ ATACTGATCGTTGAGCGAATTTTGGC 3′) and 1752 (5′ CACCAGCCTCCAAACCAG 3′).

### Verification of transformant phenotype

Strains were mated in liquid synthetic crossing media (SC) with filter paper and no added sucrose. Strains 5941 and W1432 were used as sensitive tester strains. Strains 4739 and W1446 were used as killer tester strain. These strains were inoculated into culture tubes and allowed to grow for 3 days. Conidia from knockout strains were then spread over these cultures and cultures were allowed to mate over a 2-week period. Once ascospores could first be observed on the walls of the culture tubes, mature perithecia were removed and dissected. Rosettes of ascospores were imaged at 200x magnification.

### MSUD analysis

Four *Sk-1* strains and four sensitives strains were selected and crossed to each other in an all-by-all fashion to determine the effect of genomic background on spore killing. Strains 4746, 5940, 4738 (*Sk-1*), and 4739 (*Sk-1*) were selected to represent Tahiti, and strains W1426, W1312, W1325 (*Sk-1*), and W1446 (*Sk-1*) were selected from Italy. In all cases, ascospores were randomly selected to generate progeny, conidia were harvested and used to fertilize the sensitive parent to generate backcrossed lines. For one cross each of Tahiti x Tahiti (*Sk-1*), Tahiti x Italy (*Sk-1*), Italy x Tahiti (*Sk-1*), and Italy x Italy (*Sk-1*) lines were constructed in triplicate to control for any laboratory effects.

To quantify the strength of the loss of spore killing, new crosses were conducted between all parental strains. Perithecia were harvested and dissected at maturity and asci were scored for spore killing as either: killing (4-spored), no killing (8-spored), or intermedia (4 -- 8 -spored) by manual counts under a dissecting microscope. Loss of killing was confirmed independently in two separate laboratories. The ability of F1 strains to perform spore killing when outcrossed was evaluated by crossing F1 progeny of 4746 x 4738 to the Italian sensitive strain W1426.

## End notes

### Acknowledgements

We would like to thank Katarina Bergqwist and Markus Hiltunen for help with phenotyping strains, Douglas Scofield for bioinformatics assistance and Magdalena Grudzinska-Sterno for assistance with RNA sequencing. We acknowledge support of the National Genomics Infrastructure (NGI) / Uppsala Genome Center and UPPMAX for providing assistance in massive parallel sequencing and computational infrastructure. Work performed at NGI / Uppsala Genome Center has been funded by RFI / VR and SciLifeLab, Sweden. Research performed at Illinois State University was supported by a grant from the National Science Foundation (NSF) (MCB# 1615626). The project was supported by a European Research Council grant under the program H2020, ERC-2014-CoG, project 648143 (SpoKiGen) (to HJ)

### Data and strain availability

Raw reads will be deposited to the Short Read Archive at NCBI. Genome assemblies, SNP data in VCF format and alignments will be deposited at FigShare. All *N. sitophila* strains used in this study that was previously not available from the Fungal Genetics Stock Center will be deposited there.

### Author contributions

JS and HJ planned the study. DJJ contributed with fungal material and supervision of the project. AV performed experiments on segregation distortion. NR, DS and TMH generated all mutants. JS performed genome extractions and bioinformatical analysis. JS drafted the manuscript and HJ, ML, AV, NR, and TMH contributed to the text.

## Supplementary material

**Figure S1:**
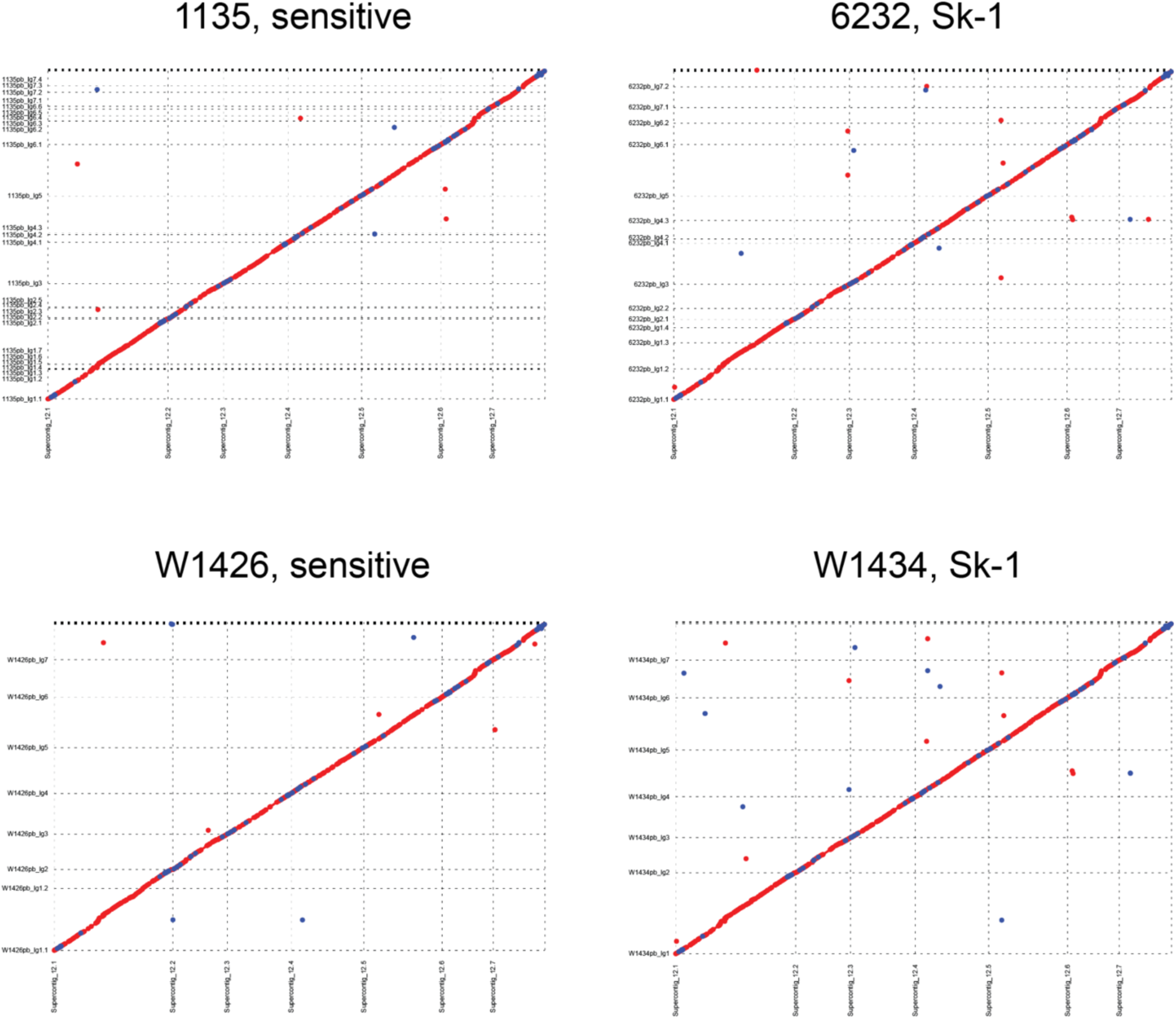
Dot plots of PacBio assemblies aligned to *N. crassa*. Dot plots showing whole genome alignment of the four PacBio assemblies of *N. sitophila* to the *N. crassa* OR74 reference assembly. PacBio contigs are plotted vertically, and *N. crassa* chromosomes horizontally.

**Figure S2:**
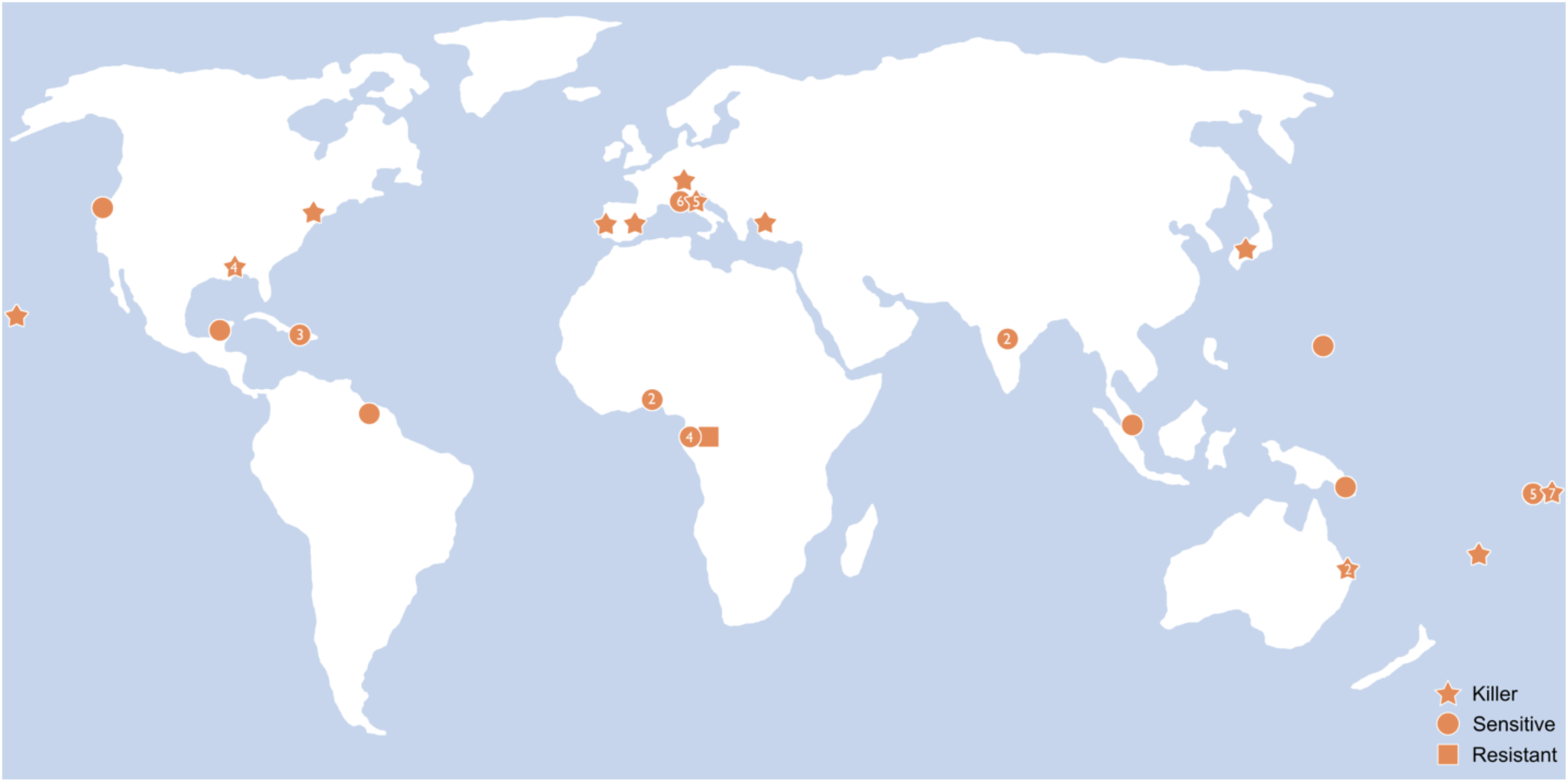
Map of samples. Map indicating sampling locations of all *N. sitophila* strains included in this study. Sensitive strains are marked with circles, *Sk-1* killers with stars and the one resistant strain with a square. Numbers in symbols represent sample-numbers at each site.

**Figure S3:**
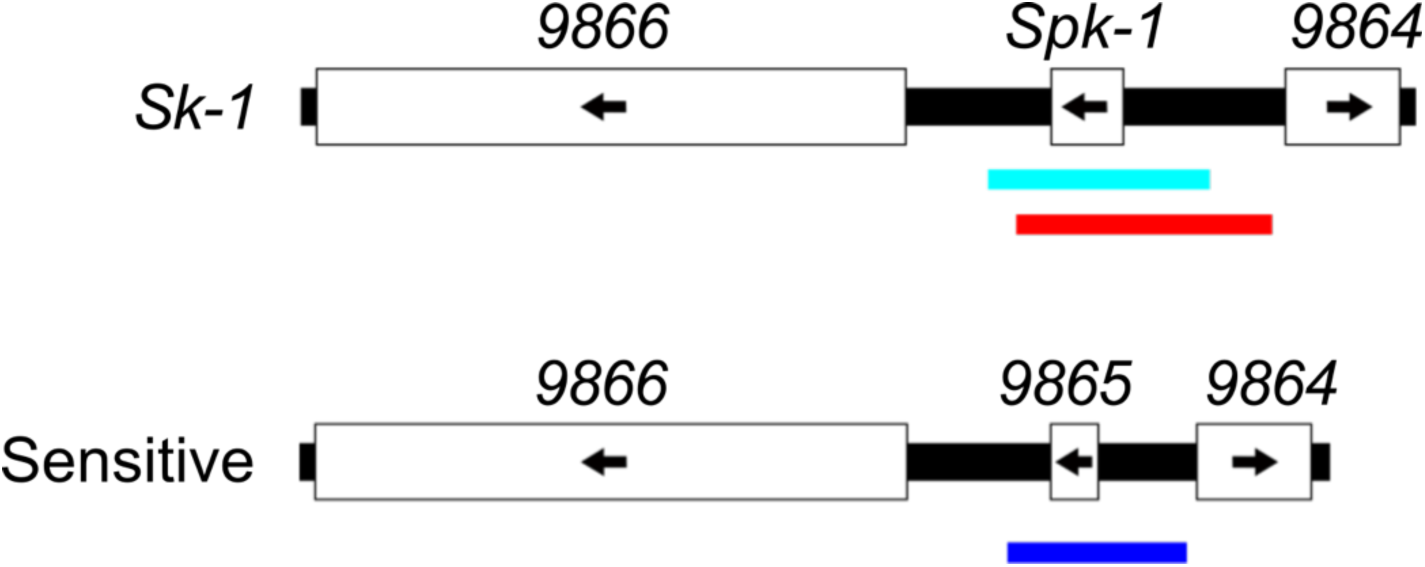
Deletion and insertion of the active element. The causative factor behind the *Sk-1 spore killing phenotype* is located between genes *NCU09866* and *NCU09864* on chromosome 6. A diagram of the *NCU09866*-*NCU09864* locus in *Sk-1* (top) strains and sensitive (bottom) strains is shown. The spore killing phenotype of an *Sk-1* strain can be eliminated by deleting a 2.8 kb interval (cyan) spanning the putative *Sk-1* gene. A spore killing phenotype can be established in a sensitive strain by replacing a 2.3 kb interval spanning *NCU0965* (blue) with the allelic 3.2 kb interval from an *Sk-1* strain (red).

**Figure S4:**
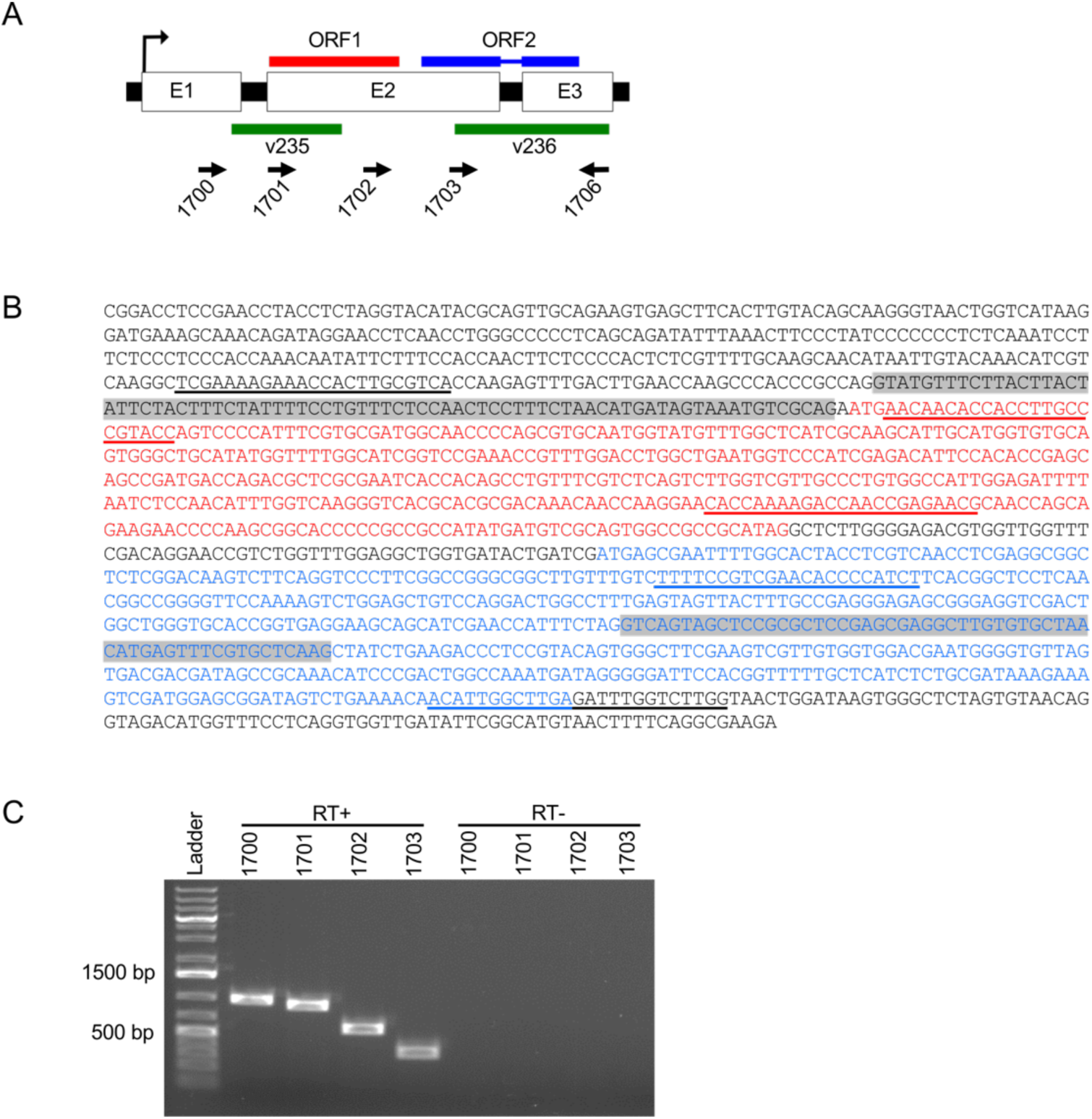
Dissection of the *Spk-1* transcript. **(A)** The *Spk-1* gene contains three exons (E1, E2, and E3) and two introns. The gene contains at least two potential open reading frames: ORF1 (red) and ORF2 (blue). The intervals deleted with vector 235 and vector 236 are indicated with green bars. The binding sites and directions of primers used in RT-PCR assays are indicated with arrows. **(B)** The *Spk-1* sequence is shown. Red and blue font are used for ORF1 and ORF2, respectively, and confirmed introns are highlighted in gray. The primer binding sites for primers 1700, 1701, 1702, 1703, and 1706 are underlined. **(C)** RT-PCR analysis was performed on total RNA from vegetative tissue of strain W1434. PCR was performed with the indicated forward primer (1700, 1701, 1702, 1703) and primer 1706 as the reverse primer. The image depicts products from each PCR reaction. Cloning and sequencing of the products in lanes “1700” and “1701” confirmed the locations of the two introns depicted in panels A and B (data not shown). No other introns were identified from the cloning and sequencing experiments. RT+, reverse transcriptase was used in the cDNA synthesis reaction; RT-, reverse transcriptase was left out of the cDNA synthesis reaction.

**Figure S5:**
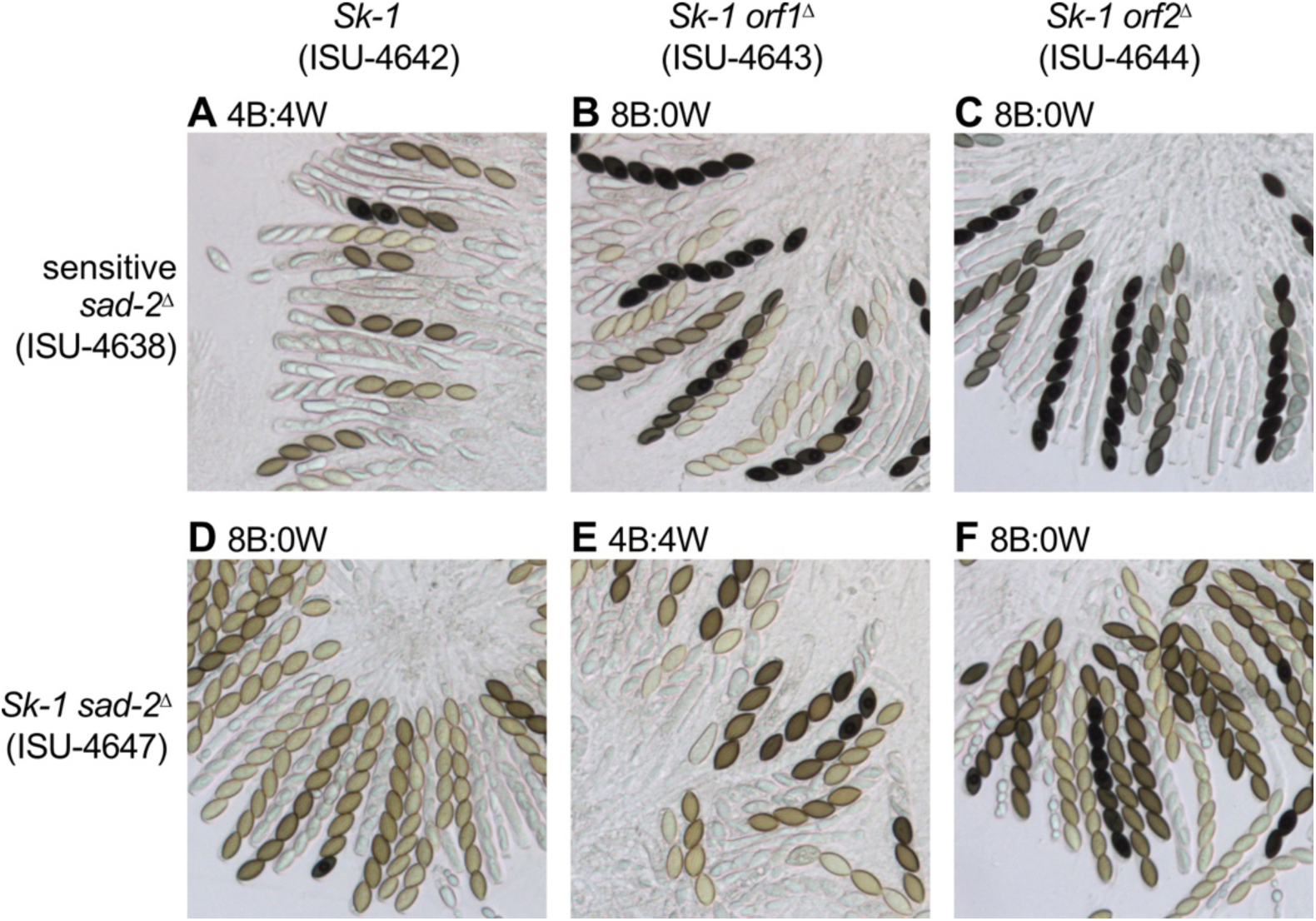
Ascus phenotypes of ORF deletions. Partial deletion of *Sk-1* ORF1 and *Sk-1* ORF2 have different effects on killing and resistance. The 5′ half of ORF1 was deleted with vector 235 to produce strain ISU-4643, and the 3′ 2/3 of ORF2 was deleted with vector 236 to produce strain ISU-4644. (A-F) Images are of asci from six crosses. The predominant phenotype is indicated above each image. The partial ORF1 deletion disrupted killing and resistance, while the partial ORF2 deletion disrupted killing but not resistance.

**Figure S6:**
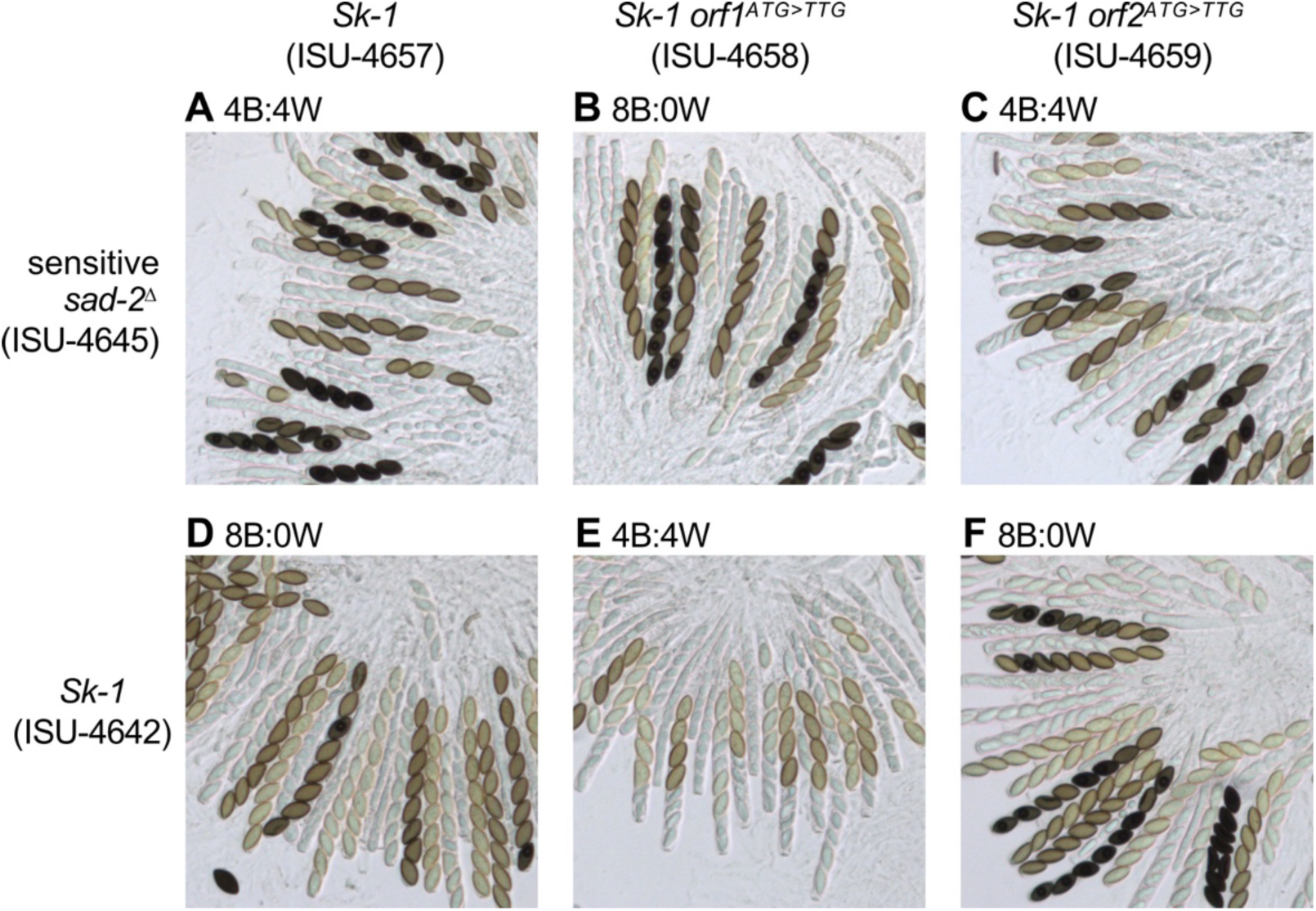
Ascus phenotypes of mutating the start codons. Mutation of the *Spk-1* ORF1 start codon eliminates spore killing and resistance to spore killing. Strains ISU-4657, ISU-4658, and ISU-4659 were generated by inserting an *Spk-1* transgene or an *Spk-1* mutant transgene at the allelic location in a sensitive strain. (A-F) Images are of asci from six crosses. The predominant phenotype is indicated above each image. Mutation of the ORF1 start codon eliminated both killing and resistance, while mutation of the ORF2 start codon had no effect on killing or resistance.

**Figure S7:**
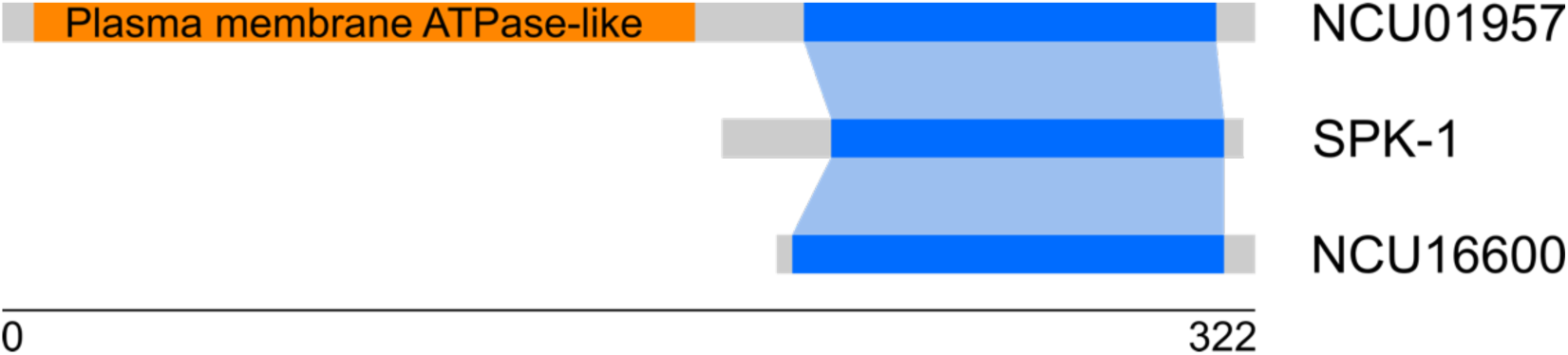
*Spk-1* shows homology to *NCU01957* and *NCU16600*. Cartoon of homologous regions of *Spk-1* and the two annotated *N. crassa* genes *NCU01957* and *NCU16600*. All three genes share the same homologous region which spans almost the entire sequence of *Spk-1* and *NCU16600*, but only the C-terminal region of the longer *NCU01957* gene. *NCU01957* also contains a region with strong similarity to *N. crassa* membrane ATPase gene, but this region is not found in the other two genes.

**Figure S8:**
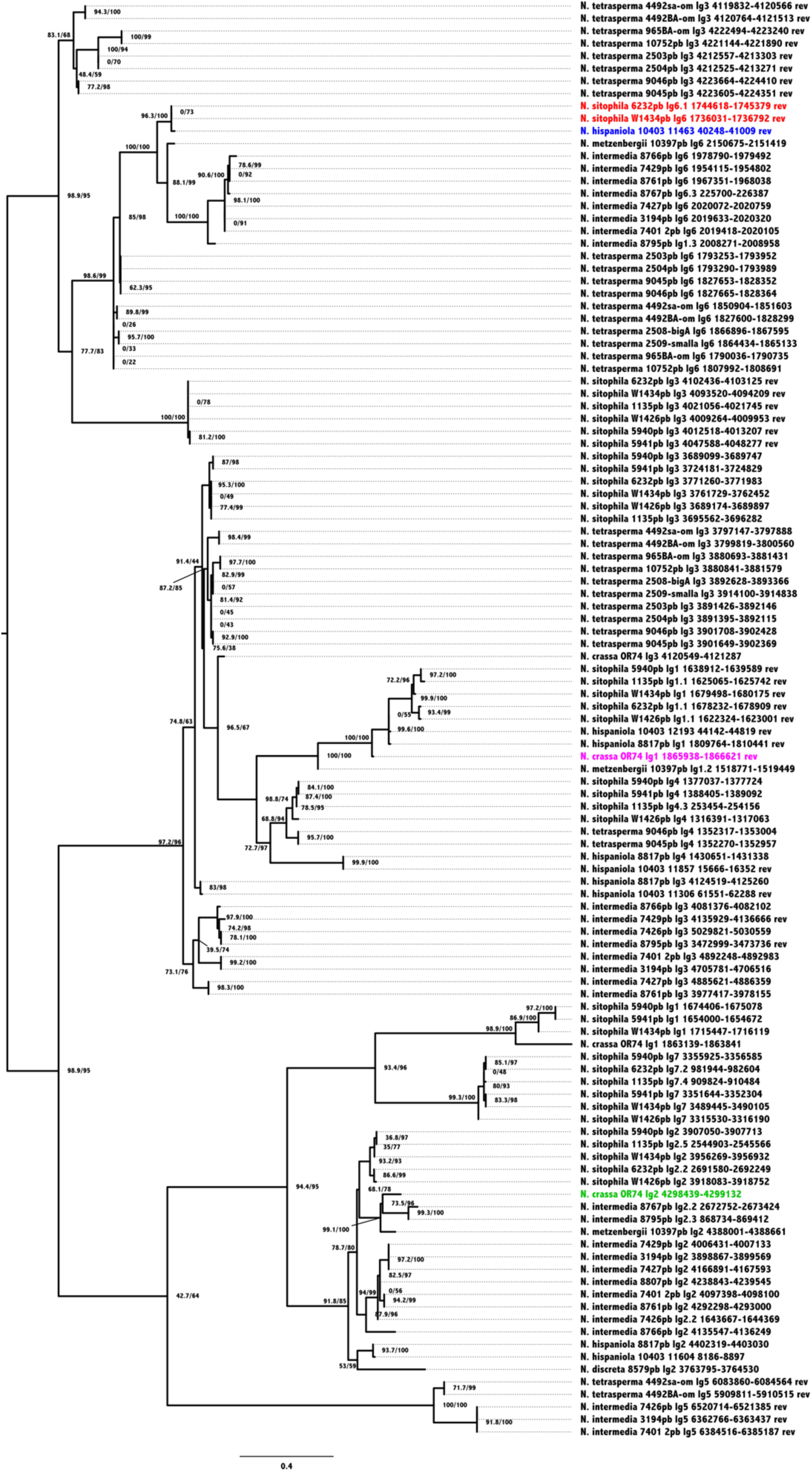
Phylogenetic tree of *Spk-1* homologs. Maximum likelihood phylogeny of SPK-1 and homologs in 31 *Neurospora* strains from 7 different species (*N. crassa*, *N. discreta*, *N. hispaniola*, *N. intermedia*, *N. metzenbergii*, *N. sitophila* and *N. tetrasperma*). SPK-1 from the two *N. sitophila Sk-1* strains are marked in red. *N. hispaniola* strain 10403 is the only other strain that carries a highly similar sequence (98.5% amino acid sequence identity, marked in blue), which is also found at the same locus on chromosome 6. Other homologous sequences range in amino acid sequence identity between 24% and 87%. Two annotated genes in the *N. crassa* OR74 reference genome are found among this set of sequences: *NCU01957* (also known as *AR2*, marked in magenta) and *NCU16600* (marked in green). *NCU01957* contains a Plasma Membrane ATPase domain not found in SPK-1 and is known to cause empty asci when mutated (Randall & Metzenberg 1998), but has otherwise no known function. *NCU16600* is also of unknown function.

**Figure S9:**
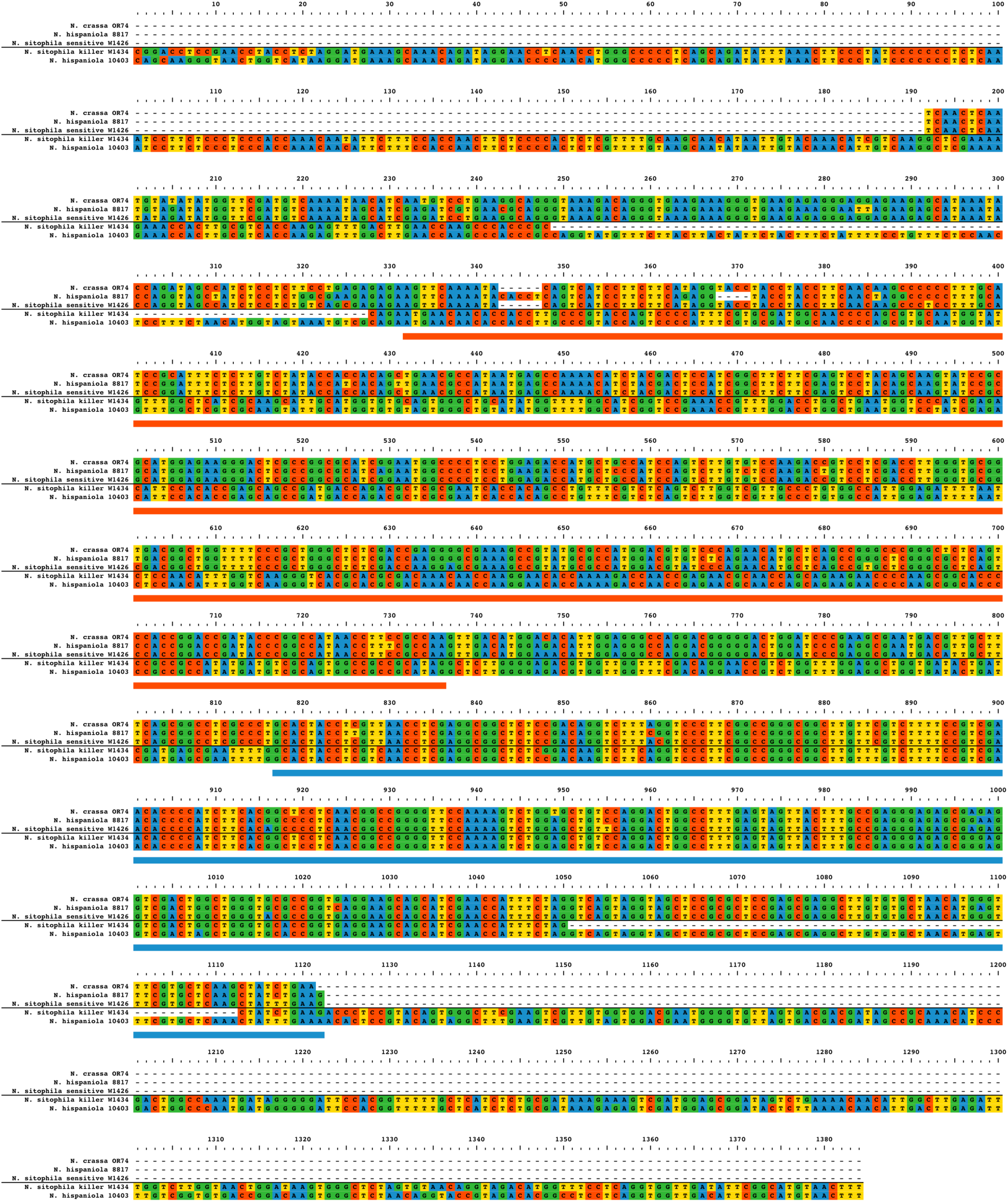
Alignment of *Spk-1* and *NCU09865* transcripts. Multiple sequence alignment of *NCU09865* from *N. crassa* OR74, *N. hispaniola* 8817 and *N. sitophila* W1426 (sensitive) and the *Spk-1* transcript from *N. sitophila* W1434 (killer) and *N. hispaniola* 10403. Only the region marked with a blue block (position 917-1122) is actually aligning between *NCU09865* and *Spk-1*, and corresponds to the truncated part of *NCU09865* that remains at the *Spk-1* locus. The red block (position 332-736) marks the open reading frame of the *Spk-1* transcript.

**Figure S10:**
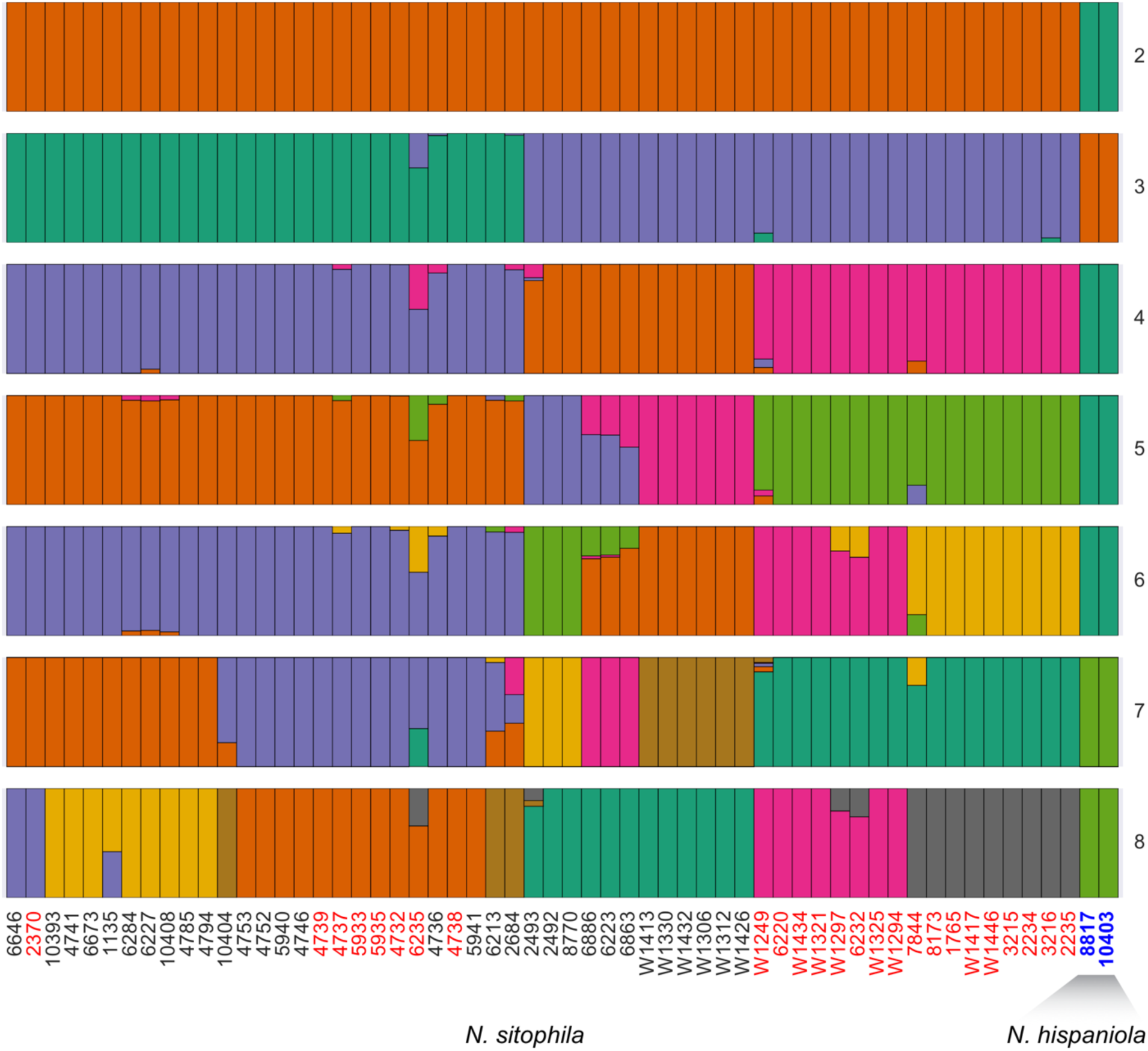
Admixture analysis of *N. sitophila* and *N. hispaniola*. An ADMIXTURE analysis of *N. sitophila* and *N. hispaniola* strains show no genome-wide signal of gene flow between the two species when varying number of clusters from 2 to 8. While *Spk-1* has been introgressed from *N. hispaniola*, geneflow must be rare.

**Figure S11:**
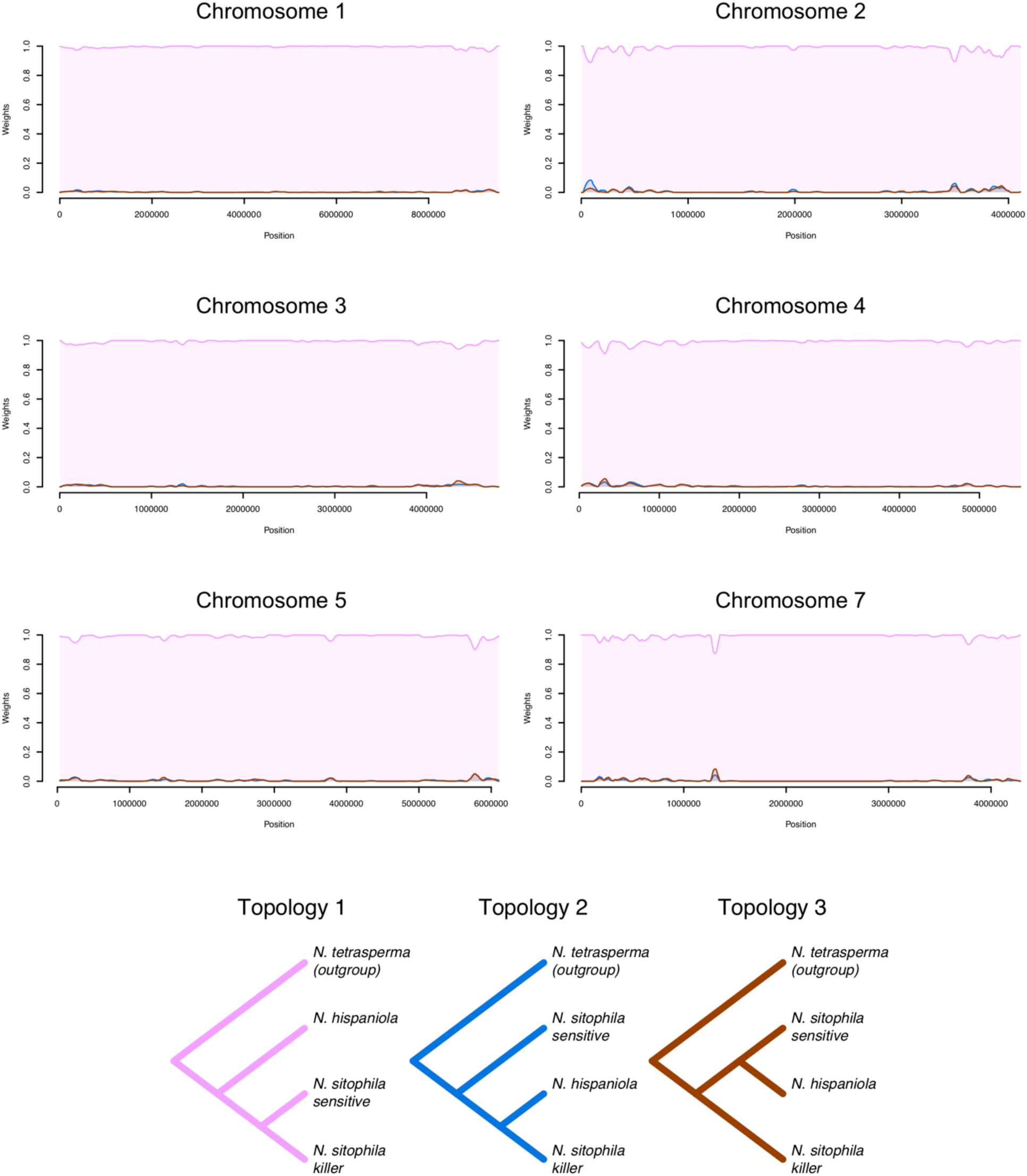
Local patterns of introgression across the genome. Twisst plots showing signals of local introgression from *N. hispaniola* across chromosome 1 to 5 and 7 (chromosome 6 is shown in Figure 2). The pink line shows the fraction of all tetrad trees that do not support introgression, while blue is consistent with introgression into *Sk-1* and brown with introgression into sensitive *N. sitophila* strains. All curves have been smoothed using Loess smoothing (span=0.05). Some small regions show a signal consistent with introgression, but no window has a stronger signal than *Spk-1*.

**Figure S12:**
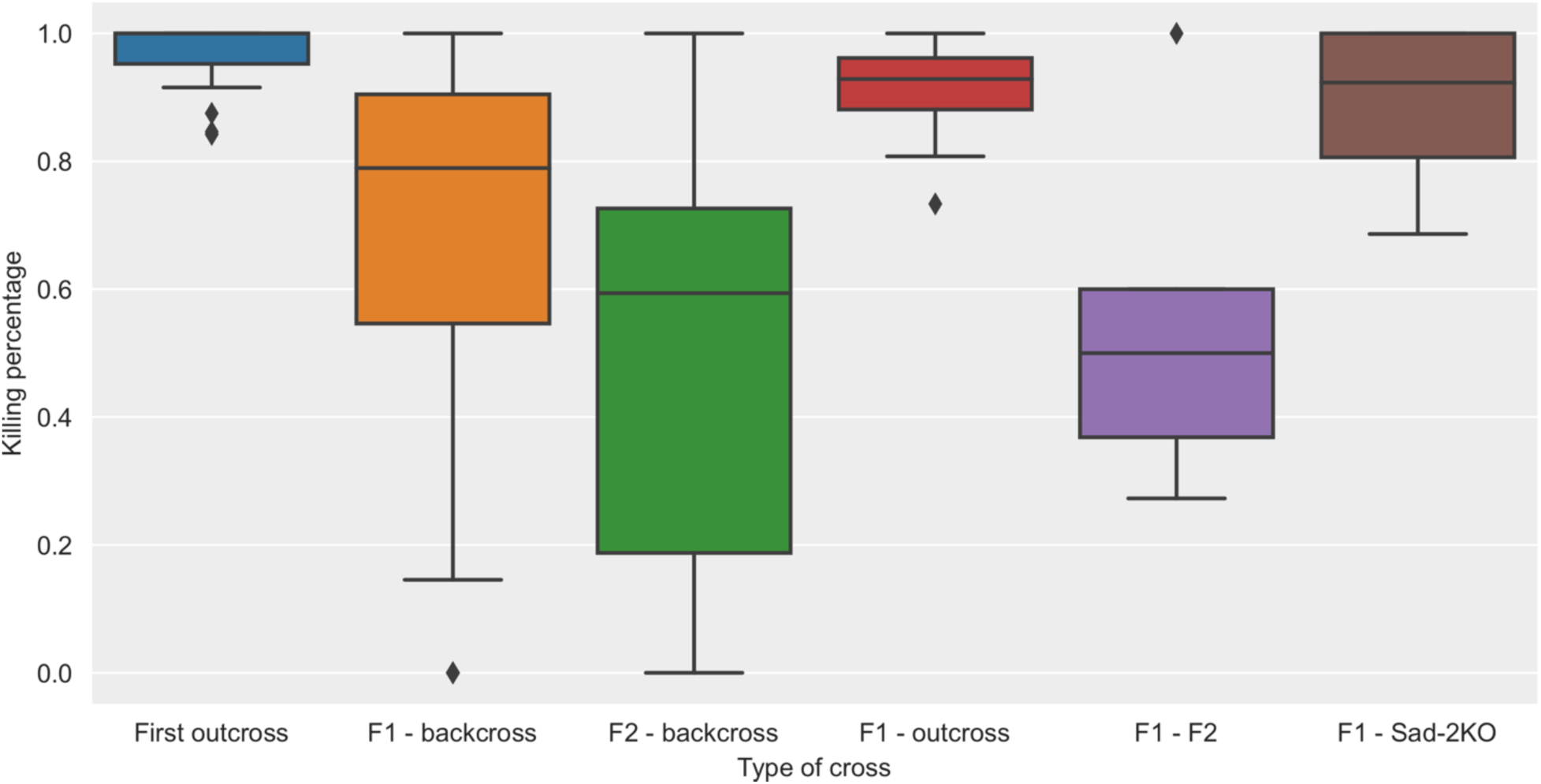
MSUD suppressed spore killing. Boxplot of fraction of asci containing 4 spores, indicating that spore killing has happened. Data is based on crosses between the sensitive Tahiti strains 4746 and 5940 and the killer strains 4738, 4739, W1325 and W1446. “Outcrosses” show the killing percentage in these crosses, “F1 - backcross” show the first backcross between the F1 and the sensitive parental strain and “F2 - backcross”, the second backcross. “F1 - outcross” shows killing percentage of an outcross between a killer F1 with a nonparental sensitive strain. “F1 - F2” shows crosses between a killer F1 and a sensitive F2. “F1 - sad-2KO” shows the killing percentage in a cross between the killer F1 and its sensitive parent, where the *sad-2* gene has been deleted. This deactivates the MSUD system.

**Figure S13:**
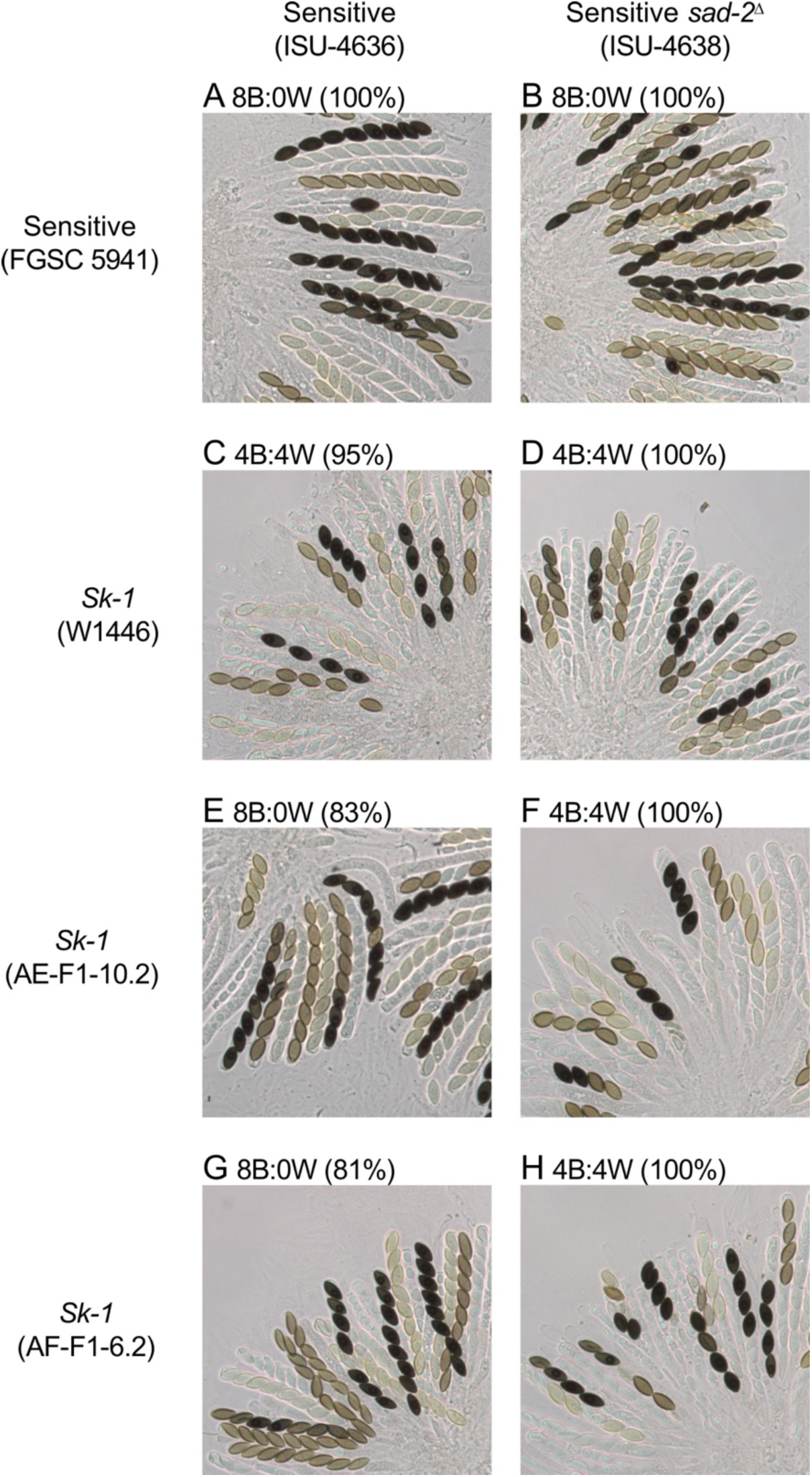
Ascus phenotype of MSUD suppressed spore killing. Suppression of MSUD increases the efficiency of spore killing in *Sk-1* × sensitive crosses. (A-H) The images are of rosettes from eight different crosses. The predominant ascus phenotype for each cross is listed above each image, along with the percentage of asci showing the predominant phenotype. AE-F1-10.2 is an F1 offspring from a cross between 4739 and 4746, and AF-F1-6.2 is an F1 offspring from a cross between 4738 and 4746.

**Table S1:** PacBio sequencing and genome assembly statistics for four N. sitophila strains

| Strain [1] | Phenotype | Location | Mean coverage | Mean subread length (bp) | Assembly size (Mbp) | Contigs | Mapped contigs [2] |
| --- | --- | --- | --- | --- | --- | --- | --- |
| 6232 | *Sk-1* | Turkey | 146 | 6038 | 39 | 27 | 15 |
| 1135 | Sensitive | Panama | 143 | 6175 | 41.2 | 42 | 20 |
| W1426 | *Sk-1* | Italy | 93.5 | 7432 | 37.9 | 20 | 8 |
| W1434 | Sensitive | Italy | 80.2 | 8197 | 39.2 | 14 | 7 |

[1] 1135 and 6232 are strain ID numbers in the Fungal Genetics Stock Center (FGSC). W1426 and W1434 are ID numbers from D. Jacobson's personal collection.
[2] Number of contigs mapping to the seven chromosomes of the *N. crassa* OR74A reference genome.

**Table S2:** Illumina HiSeq sequencing and assembly statistics

| Species | Phenotype | Location | Strain | Contigs | Assembly size (bp) | N90 | N80 | N50 | N20 | Largest contig | Number of reads | Read length | Total data (Gbp) | Coverage |
| --- | --- | --- | --- | --- | --- | --- | --- | --- | --- | --- | --- | --- | --- | --- |
| *N. sitophila* | Sk-1 | Japan | 1765 | 1953 | 38,707,971 | 15,821 | 25,744 | 55,111 | 114,458 | 371,488 | 8,612,848 | 124 | 2.14 | 55.2 |
| *N. sitophila* | Sk-1 | USA | 2234 | 1863 | 38,485,632 | 17,629 | 30,042 | 71,689 | 135,644 | 272,731 | 10,913,177 | 124 | 2.71 | 70.3 |
| *N. sitophila* | Sk-1 | USA | 2235 | 1805 | 38,772,771 | 17,922 | 29,059 | 64,944 | 121,534 | 319,446 | 12,202,102 | 124 | 3.03 | 78.0 |
| *N. sitophila* | Sk-1 | Hawaii | 2370 | 1636 | 37,890,652 | 17,190 | 26,072 | 55,709 | 112,741 | 338,630 | 10,212,922 | 124 | 2.53 | 66.8 |
| *N. sitophila* | Sensitive | India | 2492 | 1904 | 38,128,085 | 17,125 | 27,456 | 62,599 | 121,274 | 448,962 | 12,570,109 | 124 | 3.12 | 81.8 |
| *N. sitophila* | Sensitive | Malaya | 2493 | 1747 | 38,881,544 | 18,233 | 28,089 | 66,516 | 122,833 | 351,117 | 11,389,221 | 124 | 2.82 | 72.6 |
| *N. sitophila* | Sk-1 | USA | 2684 | 1371 | 38,159,651 | 21,773 | 33,360 | 74,598 | 145,207 | 950,644 | 10,429,030 | 124 | 2.59 | 67.8 |
| *N. sitophila* | Sk-1 | USA | 3215 | 1730 | 38,297,183 | 18,010 | 28,739 | 69,997 | 140,229 | 295,300 | 9,679,280 | 124 | 2.40 | 62.7 |
| *N. sitophila* | Sk-1 | USA | 3216 | 1828 | 38,623,104 | 17,025 | 28,503 | 60,409 | 119,478 | 304,077 | 9,465,706 | 124 | 2.35 | 60.8 |
| *N. sitophila* | Sk-1 | Tahiti | 4732 | 1591 | 38,167,904 | 16,834 | 27,134 | 60,676 | 116,640 | 379,184 | 10,047,815 | 124 | 2.49 | 65.3 |
| *N. sitophila* | Sensitive | Tahiti | 4736 | 1906 | 38,155,269 | 18,038 | 28,819 | 63,253 | 125,960 | 396,913 | 14,128,440 | 124 | 3.50 | 91.8 |
| *N. sitophila* | Sk-1 | Tahiti | 4737 | 1705 | 38,289,076 | 16,369 | 26,517 | 60,017 | 116,911 | 389,253 | 10,723,121 | 124 | 2.66 | 69.5 |
| *N. sitophila* | Sk-1 | Tahiti | 4738 | 2081 | 37,961,582 | 16,009 | 26,042 | 56,517 | 114,157 | 379,162 | 11,263,599 | 124 | 2.79 | 73.6 |
| *N. sitophila* | Sk-1 | Tahiti | 4739 | 1637 | 38,083,069 | 15,983 | 25,694 | 55,427 | 117,519 | 397,723 | 10,136,193 | 124 | 2.51 | 66.0 |
| *N. sitophila* | Sensitive | Papua New Guinea | 4741 | 2176 | 38,276,791 | 16,370 | 25,598 | 53,346 | 111,183 | 326,143 | 9,662,540 | 124 | 2.40 | 62.6 |
| *N. sitophila* | Sensitive | Tahiti | 4746 | 1915 | 37,979,133 | 15,506 | 25,547 | 55,239 | 114,037 | 255,821 | 9,632,767 | 124 | 2.39 | 62.9 |
| *N. sitophila* | Sensitive | Tahiti | 4752 | 2769 | 38,447,232 | 16,083 | 25,038 | 55,913 | 95,867 | 249,085 | 10,092,165 | 124 | 2.50 | 65.1 |
| *N. sitophila* | Sensitive | Tahiti | 4753 | 1700 | 37,981,914 | 17,728 | 27,775 | 60,433 | 118,535 | 414,591 | 10,960,501 | 124 | 2.72 | 71.6 |
| *N. sitophila* | Sensitive | Haiti | 4785 | 1622 | 37,896,451 | 16,375 | 23,869 | 53,142 | 103,698 | 396,448 | 10,313,457 | 124 | 2.56 | 67.5 |
| *N. sitophila* | Sensitive | Haiti | 4794 | 1462 | 37,917,033 | 18,260 | 28,253 | 64,044 | 122,270 | 414,635 | 10,455,431 | 124 | 2.59 | 68.4 |
| *N. sitophila* | Sk-1 | Tahiti | 5933 | 2078 | 38,048,338 | 16,864 | 26,397 | 57,685 | 110,222 | 316,643 | 9,675,980 | 124 | 2.40 | 63.1 |
| *N. sitophila* | Sk-1 | Tahiti | 5935 | 1923 | 38,151,115 | 16,972 | 26,217 | 55,129 | 115,663 | 389,311 | 9,566,368 | 124 | 2.37 | 62.2 |
| *N. sitophila* | Sensitive | Tahiti | 5940 | 2067 | 37,864,153 | 17,040 | 26,066 | 54,872 | 106,647 | 349,123 | 11,339,082 | 124 | 2.81 | 74.3 |
| *N. sitophila* | Sensitive | Tahiti | 5941 | 1701 | 37,867,531 | 17,078 | 27,426 | 60,198 | 130,267 | 262,542 | 10,771,936 | 124 | 2.67 | 70.5 |
| *N. sitophila* | Sensitive | Rota | 6213 | 1648 | 37,813,855 | 16,908 | 28,778 | 63,418 | 124,490 | 451,800 | 10,210,315 | 124 | 2.53 | 67.0 |
| *N. sitophila* | Sk-1 | USA | 6220 | 1710 | 39,377,771 | 17,989 | 29,191 | 65,081 | 130,637 | 365,571 | 11,578,623 | 124 | 2.87 | 72.9 |
| *N. sitophila* | Sensitive | Gabon | 6223 | 1951 | 37,725,619 | 17,358 | 28,384 | 58,805 | 108,770 | 282,389 | 9,537,071 | 124 | 2.37 | 62.7 |
| *N. sitophila* | Sensitive | Gabon | 6227 | 3019 | 38,139,207 | 13,525 | 21,277 | 45,411 | 85,486 | 375,307 | 7,681,545 | 124 | 1.91 | 49.9 |
| *N. sitophila* | Sk-1 | Turkey | 6232 | 2106 | 38,898,329 | 16,791 | 26,063 | 61,467 | 123,557 | 464,739 | 9,737,681 | 124 | 2.41 | 62.1 |
| *N. sitophila* | Sk-1 | Roratonga | 6235 | 2070 | 38,313,666 | 15,580 | 25,215 | 56,854 | 115,919 | 282,713 | 9,119,530 | 124 | 2.26 | 59.0 |
| *N. sitophila* | Sensitive | Ivory coast | 6284 | 1549 | 37,850,405 | 17,836 | 28,310 | 60,190 | 121,625 | 471,273 | 9,509,623 | 124 | 2.36 | 62.3 |
| *N. sitophila* | Sensitive | Mexico | 6646 | 1548 | 37,917,104 | 17,406 | 27,952 | 61,709 | 113,577 | 413,832 | 9,655,422 | 124 | 2.39 | 63.2 |
| *N. sitophila* | Sensitive | Brazil | 6673 | 1618 | 37,932,419 | 16,753 | 26,308 | 54,172 | 108,750 | 262,918 | 8,904,810 | 124 | 2.21 | 58.2 |
| *N. sitophila* | Sensitive | Hawaii | 6689 | 10327 | 43,985,540 | 11,677 | 24,931 | 62,163 | 121,934 | 383,175 | 9,312,497 | 124 | 2.31 | 52.5 |
| *N. sitophila* | Resistant | Gabon | 6850 | 1629 | 39,107,064 | 16,857 | 28,995 | 60,589 | 111,166 | 290,727 | 9,339,131 | 124 | 2.32 | 59.2 |
| *N. sitophila* | Sensitive | Ivory coast | 6863 | 1645 | 37,789,059 | 17,514 | 28,023 | 59,451 | 114,104 | 283,786 | 9,865,808 | 124 | 2.45 | 64.7 |
| *N. sitophila* | Sensitive | Gabon | 6886 | 1892 | 37,863,691 | 16,625 | 26,786 | 60,446 | 114,693 | 378,496 | 9,039,032 | 124 | 2.24 | 59.2 |
| *N. sitophila* | Sk-1 | Australia | 7844 | 1884 | 38,762,514 | 16,500 | 26,795 | 57,191 | 107,482 | 409,024 | 9,172,136 | 124 | 2.27 | 58.7 |
| *N. sitophila* | Sk-1 | Australia | 8173 | 1834 | 38,493,069 | 17,297 | 26,291 | 59,376 | 119,284 | 279,482 | 9,896,780 | 124 | 2.45 | 63.8 |
| *N. sitophila* | Sensitive | India | 8770 | 1865 | 38,113,580 | 16,423 | 25,764 | 56,769 | 112,688 | 319,637 | 9,972,863 | 124 | 2.47 | 64.9 |
| *N. sitophila* | Sensitive | Papua New Guinea | 10393 | 1491 | 37,800,550 | 18,002 | 26,831 | 56,392 | 112,475 | 395,346 | 9,255,830 | 124 | 2.30 | 60.7 |
| *N. sitophila* | Sensitive | Haiti | 10404 | 1813 | 37,873,810 | 16,452 | 26,738 | 54,719 | 102,744 | 233,791 | 9,910,163 | 124 | 2.46 | 64.9 |
| *N. sitophila* | Sensitive | Gabon | 10408 | 1887 | 38,062,587 | 18,152 | 30,476 | 66,469 | 112,349 | 471,331 | 11,287,709 | 124 | 2.80 | 73.5 |
| *N. sitophila* | *Sk-1* | Portugal | W1249 | 1465 | 39,251,111 | 16,537 | 26,375 | 57,314 | 110,291 | 363,664 | 9,533,549 | 124 | 2.36 | 60.2 |
| *N. sitophila* | *Sk-1* | Spain | W1294 | 2329 | 39,072,114 | 15,077 | 23,893 | 51,772 | 102,729 | 494,286 | 10,188,425 | 124 | 2.53 | 64.7 |
| *N. sitophila* | *Sk-1* | Switzerland | W1297 | 1930 | 39,216,768 | 16,185 | 25,216 | 56,776 | 107,778 | 554,470 | 9,694,420 | 124 | 2.40 | 61.3 |
| *N. sitophila* | Sensitive | Italy | W1306 | 1828 | 37,708,235 | 16,451 | 26,515 | 57,583 | 99,977 | 296,869 | 9,734,942 | 124 | 2.41 | 64.0 |
| *N. sitophila* | Sensitive | Italy | W1312 | 1821 | 37,699,980 | 16,753 | 27,118 | 61,471 | 113,498 | 425,044 | 10,853,639 | 124 | 2.69 | 71.4 |
| *N. sitophila* | *Sk-1* | Italy | W1321 | 2069 | 39,410,520 | 15,969 | 25,193 | 58,514 | 110,590 | 420,290 | 9,682,315 | 124 | 2.40 | 60.9 |
| *N. sitophila* | *Sk-1* | Italy | W1322 | 1883 | 37,188,009 | 17,387 | 28,626 | 59,733 | 116,387 | 306,637 | 9,978,456 | 124 | 2.47 | 66.5 |
| *N. sitophila* | *Sk-1* | Italy | W1325 | 2175 | 39,108,556 | 16,752 | 26,205 | 57,541 | 112,927 | 321,839 | 8,882,548 | 124 | 2.20 | 56.3 |
| *N. sitophila* | Sensitive | Italy | W1330 | 1827 | 37,733,606 | 15,518 | 25,334 | 57,073 | 99,718 | 424,769 | 10,153,224 | 124 | 2.52 | 66.7 |
| *N. sitophila* | Sensitive | Italy | W1413 | 2114 | 37,600,533 | 12,686 | 20,141 | 41,628 | 87,404 | 293,508 | 9,301,378 | 124 | 2.31 | 61.3 |
| *N. sitophila* | *Sk-1* | Italy | W1417 | 1805 | 38,625,776 | 16,389 | 26,201 | 60,780 | 116,147 | 381,804 | 11,101,624 | 124 | 2.75 | 71.3 |
| *N. sitophila* | Sensitive | Italy | W1426 | 2018 | 37,649,517 | 13,626 | 21,600 | 45,376 | 88,860 | 293,454 | 7,766,246 | 124 | 1.93 | 51.2 |
| *N. sitophila* | Sensitive | Italy | W1432 | 2040 | 37,712,739 | 13,481 | 22,012 | 45,587 | 90,778 | 293,510 | 7,537,534 | 124 | 1.87 | 49.6 |
| *N. sitophila* | *Sk-1* | Italy | W1434 | 2258 | 38,831,428 | 12,628 | 20,340 | 41,441 | 88,905 | 300,902 | 7,398,586 | 124 | 1.83 | 47.3 |
| *N. sitophila* | *Sk-1* | Italy | W1446 | 2036 | 38,623,459 | 14,092 | 22,116 | 51,420 | 104,107 | 598,164 | 7,793,787 | 124 | 1.93 | 50.0 |
| *N. hispaniola* | Unknown | Haiti | 10403 | 2154 | 41,172,793 | 22,332 | 34,629 | 71,246 | 132,013 | 433,467 | 10,136,566 | 124 | 2.51 | 61.1 |
| *N. perkinsii* | Sensitive | Congo | 8842 | 2266 | 42,659,328 | 20,719 | 35,046 | 74,666 | 147,897 | 327,068 | 9,702,211 | 124 | 2.41 | 56.4 |
| *N. perkinsii* | Unknown | Gabon | 10406 | 2089 | 43,673,595 | 21,438 | 35,654 | 74,160 | 140,814 | 427,677 | 10,105,447 | 124 | 2.51 | 57.4 |

**Table S3:** Phenotype of *Spork1* dissection

**Table B** Phenotype summary
| Phenotype<sup>a</sup> | *Sk-1* | *Sk-1 orf1*<sup>Δ</sup> | *Sk-1 orf1*<sup>ATG>TTG</sup> | *Sk-1 orf2*<sup>Δ</sup> | *Sk-1 orf2*<sup>ATG>TTG</sup> |
| --- | --- | --- | --- | --- | --- |
| Killer | + | — | — | — | + |
| Resistant | + | — | — | + | + |

<sup>a</sup>killer: (+) The predominant ascus phenotype in crosses of the indicated genotype with sensitive is 4B:4W. resistant: (+) The predominant ascus phenotype in crosses of the indicated genotype with *Sk-1* is 8B:0W.

**Table S4:** Spork1 homologs in Neurospora

| **Species** | **Strain** | **Number of homologs [1]** |
| --- | --- | --- |
| *N. crassa* | OR74 | 4 |
| *N. discreta* | 8579 | 1 |
| *N. hispaniola* | 10403 | 5 |
| *N. hispaniola* | 8817 | 4 |
| *N. intermedia* | 3194 | 4 |
| *N. intermedia* | 7401 | 4 |
| *N. intermedia* | 7426 | 3 |
| *N. intermedia* | 7427 | 3 |
| *N. intermedia* | 7429 | 3 |
| *N. intermedia* | 8761 | 3 |
| *N. intermedia* | 8766 | 3 |
| *N. intermedia* | 8767 | 2 |
| *N. intermedia* | 8795 | 3 |
| *N. intermedia* | 8807 | 1 |
| *N. metzenbergii* | 10397 | 3 |
| *N. sitophila* | 1135 | 6 |
| *N. sitophila* | 5940 | 6 |
| *N. sitophila* | 5941 | 5 |
| *N. sitophila* | 6232 | 6 |
| *N. sitophila* | W1426 | 6 |
| *N. sitophila* | W1434 | 7 |
| *N. tetrasperma* | 2508 | 2 |
| *N. tetrasperma* | 2509 | 2 |
| *N. tetrasperma* | 4492A | 4 |
| *N. tetrasperma* | 4492a | 4 |
| *N. tetrasperma* | 965A | 3 |
| *N. tetrasperma* | 10752 | 3 |
| *N. tetrasperma* | 2503 | 3 |
| *N. tetrasperma* | 2504 | 3 |
| *N. tetrasperma* | 9045 | 4 |
| *N. tetrasperma* | 9046 | 4 |

[1] Number of hits longer than 100 aa, when searching genome assembly using tblastn.

**Table S5:** Tester strains for phenotyping spore killing in natural *N. sitophila* isolates

| **FGSC number [1]** | **Mating type** | **Phenotype** |
| --- | --- | --- |
| 4762 | A | Killer |
| 4763 | a | Killer |
| 4887 | A | Sensitive |
| 4888 | a | Sensitive |

All tester strains carry the *fluffy* mutation, which increases fertility and reduces conidiation.
[1] Strain ID numbers at the Fungal Genetics Stock Center.

**Table S6:** N. sitophila tester strains

| Name (alias) | Genotype |
| --- | --- |
| AF-F1-6.2 | *Sk-1 a* (Recombinant from FGSC 4739 × FGSC 4746) |
| AE-F1-10.2 | *Sk-1 a* (Recombinant from FGSC 4738 × FGSC 4746) |
| FGSC 4738 | *Sk-1 a* (Tahiti, 1983) |
| FGSC 4739 | *Sk-1 a* (Tahiti, 1983) |
| FGSC 4746 | *A* (Tahiti) |
| FGSC 5941 | *a* (Tahiti, 1983) |
| ISU-4296 (HDS45.4.2) | *Sk-1<sup>Δ</sup>::hph A* (Transformation of W1434 with vector 134) |
| ISU-4529 (HNR143.2.1) | *mus-51<sup>Δ</sup>::nat1; ncu09865<sup>Δ</sup>::Sk-1-hph A* (Transformation of ISU-4637 with vector 205a) |
| ISU-4636<sup>a</sup> (HNR101.14.11) | *mus-51<sup>Δ</sup>::nat1 A* (Deletion of *mus-51* from FGSC 4746) |
| ISU-4637<sup>a</sup> (HNR126.14.1) | *mus-51<sup>Δ</sup>::nat1 A* (Deletion of *mus-51* from W1426) |
| ISU-4638 (HNR119.1.1) | *mus-51<sup>Δ</sup>::nat1; sad-2<sup>Δ</sup>::hph; A* (Transformation of ISU-4636 with vector 178). |
| ISU-4641 (RNR167.28) | *mus-51<sup>Δ</sup>::nat1 a* (Recombinant from ISU-4638 × FGSC 5941) |
| ISU-4642 (RNR173.5) | *mus-51<sup>Δ</sup>::nat1; Sk-1 a* (Recombinant from W1434 × ISU-4641) |
| ISU-4643 (HNR176.1.1) | *mus-51<sup>Δ</sup>::nat1; Sk-1 orf1<sup>Δ</sup>::hph a* (Transformation of ISU-4642 with v235) |
| ISU-4644 (HNR177.1.1) | *mus-51<sup>Δ</sup>::nat1; Sk-1 orf2<sup>Δ</sup>::hph a* (Transformation of ISU-4642 with v236) |
| ISU-4645 (HNR167.17) | *mus-51<sup>Δ</sup>::nat1; sad-2<sup>Δ</sup>::hph a* (Recombinant from ISU-4638 × FGSC 5941) |
| ISU-4646 (RNR173.2) | *mus-51<sup>Δ</sup>::nat1; Sk-1 A* (Recombinant from W1434 × ISU-4641) |
| ISU-4647 (RNR226.4) | *mus-51<sup>Δ</sup>::nat1; sad-2<sup>Δ</sup>::hph; Sk-1 A* (Recombinant from ISU-4638 × W1434) |
| ISU-4657 (HNR222.1.1) | *mus-51<sup>Δ</sup>::nat1; ncu09865<sup>Δ</sup>::Sk-1-hph A* (Transformation of ISU-4636 with v205a) |
| ISU-4658 (HNR223.1.1) | *mus-51<sup>Δ</sup>::nat1; ncu09865<sup>Δ</sup>::Sk-1<sup>ORF1[ATG>TTG]</sup>-hph A* (Transformation of ISU-4636 with v205b) |
| ISU-4659 (HNR224.3.2) | *mus-51<sup>Δ</sup>::nat1; ncu09865<sup>Δ</sup>::Sk-1<sup>ORF2[ATG>TTG]</sup>-hph A* (Transformation of ISU-4636 with v205c) |
| W1426 | *A* (Italy) |
| W1434 | *Sk-1 A* (Italy) |
| W1446 | *Sk-1 a* (Italy) |

<sup>a</sup>Construction of ISU-4636 and ISU-4637 is described in Rhoades et al. (https://www.biorxiv.org/content/10.1101/761130v1).

**Table S7.**
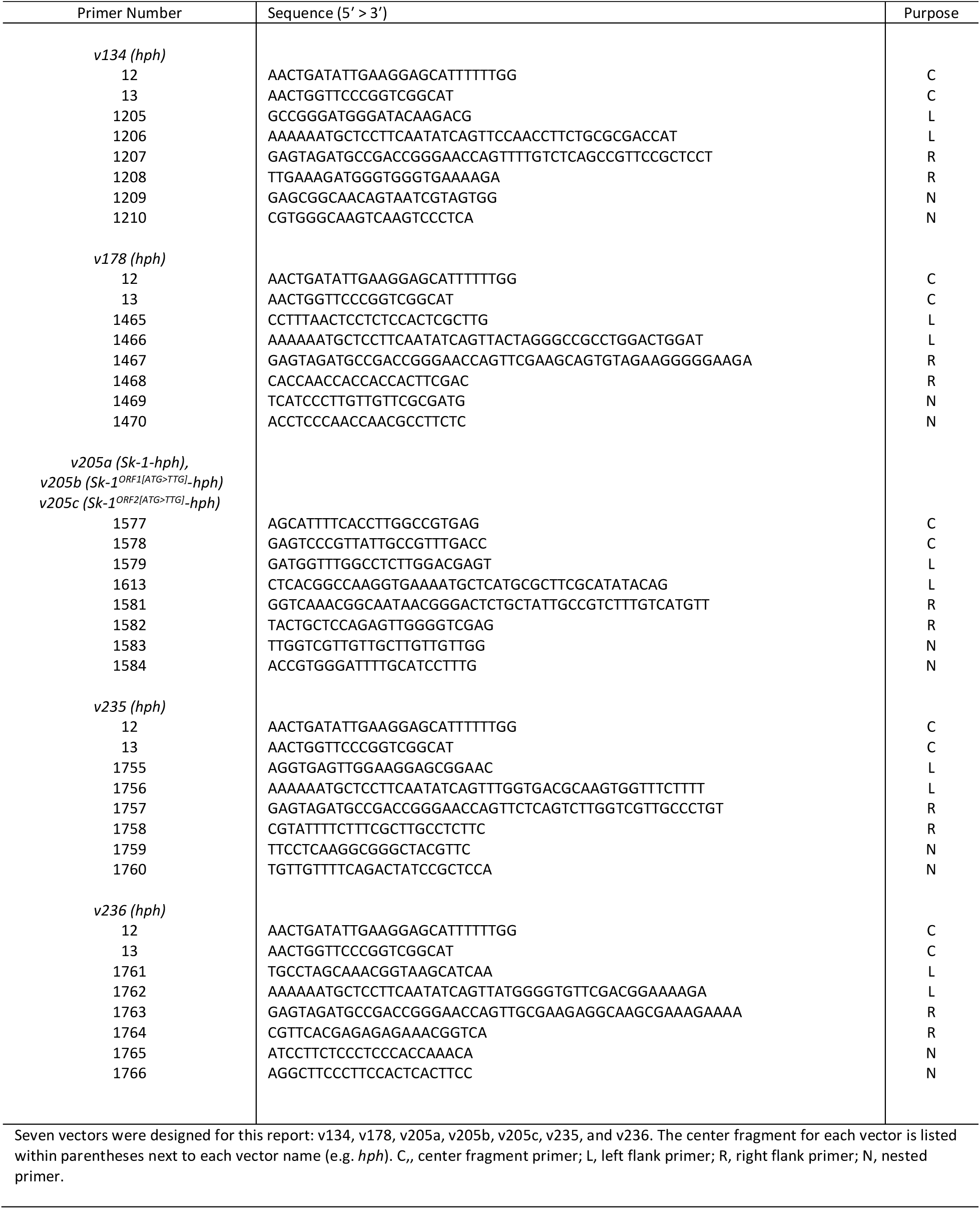
Vector construction details

